# Extending the small molecule similarity principle to all levels of biology

**DOI:** 10.1101/745703

**Authors:** Miquel Duran-Frigola, Eduardo Pauls, Oriol Guitart-Pla, Martino Bertoni, Víctor Alcalde, David Amat, Teresa Juan-Blanco, Patrick Aloy

**Affiliations:** Joint IRB-BSC-CRG Program in Computational Biology, Institute for Research in Biomedicine (IRB Barcelona), The Barcelona Institute of Science and Technology, Barcelona, Catalonia, Spain; Institució Catalana de Recerca i Estudis Avançats (ICREA), Barcelona, Catalonia, Spain

**Keywords:** Bioactivity signatures, chemical space, compound similarity principle, systems pharmacology

## Abstract

We present the Chemical Checker (CC), a resource that provides processed, harmonized and integrated bioactivity data on 800,000 small molecules. The CC divides data into five levels of increasing complexity, ranging from the chemical properties of compounds to their clinical outcomes. In between, it considers targets, off-targets, perturbed biological networks and several cell-based assays such as gene expression, growth inhibition and morphological profilings. In the CC, bioactivity data are expressed in a vector format, which naturally extends the notion of chemical similarity between compounds to similarities between bioactivity signatures of different kinds. We show how CC signatures can boost the performance of drug discovery tasks that typically capitalize on chemical descriptors, including target identification and library characterization. Moreover, we demonstrate and experimentally validate that CC signatures can be used to reverse and mimic biological signatures of disease models and genetic perturbations, options that are otherwise impossible using chemical information alone.

---

Small molecules have received a limited attention by academic researchers during the ‘omics revolution’^1^. With the exception of metabolomics, omics-based biomedical research is acutely gene-centric and difficult to link to chemical compounds^2^. The catalogue of purchasable compounds amounts to a hundred million^3^, and databases devoted to gathering bioactivity data contain a few million of them^4, 5^. Querying compounds in these databases differs greatly from querying proteins or genes, as small molecules cannot be expressed as sequence data and spell a much more complicated code. Often, the only way to characterize a small molecule is to assume it will have the same activity as compounds with similar chemical structure^2, 6^. The so-called ‘similarity principle’ has been the driving force of drug discovery. The majority of known drugs were inspired by natural products^7, 8^, chemical libraries are created by combining or decorating privileged chemotypes^9^, and the design of lead drug candidates departs from hit compounds identified in experimental screening assays^10^. Thus, compound similarities are the primary measure to chart and exploit the chemical space.

The public release of compound databases has led to the realization that the similarity principle applies beyond chemical properties. For instance, molecules with similar cell-sensitivity profiles tend to share the mechanism of action^11^, as do drugs eliciting similar side effects^12^, even when their chemical structures are unrelated. Hence, ‘biological’ similarities offer an alternative means to functionally characterize small molecules, potentially to a degree that is closer to clinical observations and beyond the mere inspection of chemical analogs^13^. Unfortunately, though, there is no convention to compare the biological profiles of small molecules, since available bioactivity data are sparse, incomplete and often of dubious quality^14^. As a result, the extent to which the similarity principle can be generalized to biology (and possibly embrace omics techniques) remains unclear. In this article, we present the Chemical Checker (CC), a resource that expands the similarity principle along the drug discovery pipeline—from *in vitro* assays to clinical observations—by treating bioactivity data with a unified analytical framework.

## Results

### Five levels of complexity for small molecule data

Of all existing compounds, approved drugs are probably the most widely characterized^15^. The way we think of drug activity suggests that small molecule data can be organized in five levels of increasing complexity (Figure 1A). A drug is often an organic molecule (A: Chemistry) that interacts with one or several protein receptors (B: Targets), triggering perturbations of biological pathways (C: Networks) and eliciting phenotypic outcomes that can be measured in e.g. cell-based assays (D: Cells) before delivery to patients (E: Clinics). Using these five categories, we classified the information stored in major compound databases, including chemogenomics resources, cell-based screens and, when available, clinical reports of drug effects (Methods).

**Figure 1.**
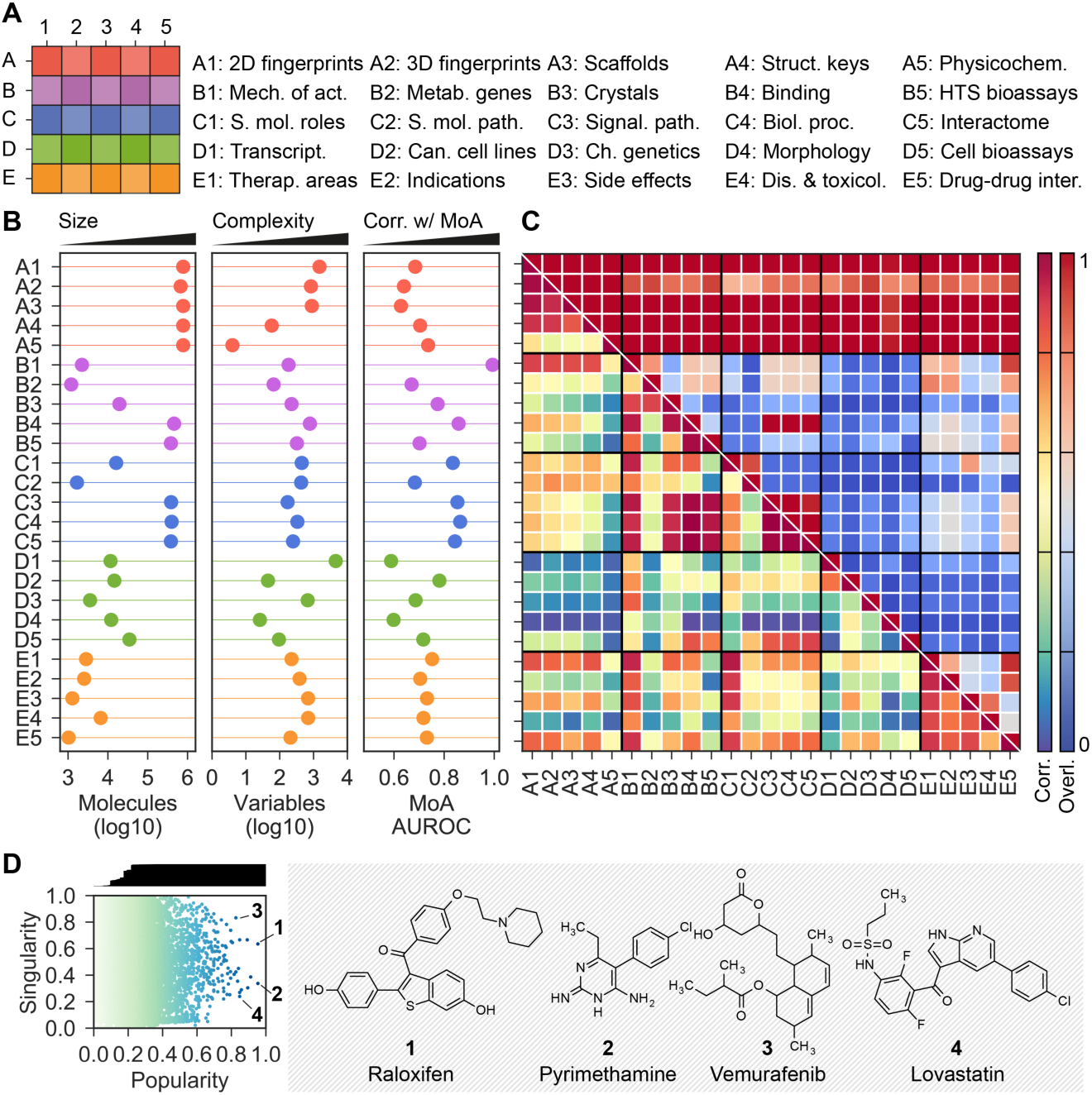
CC statistics. (A) The organization of the 5×5 CC spaces. (B) Number of molecules (size), signature length (i.e. number of latent variables as a measure of data complexity) and AUROC performances when checking if similar molecules in each CC space tend to share mechanism of action. (C) Overlap between CC spaces, in terms of number of shared molecules (upper triangle) and correlation *k* between CC spaces (lower triangle). (D) Popularity and singularity of molecules. Popularity refers to the proportion of CC spaces in which the molecule is present (correcting for correlation between CC spaces), and singularity refers to the ‘uniqueness’ of the molecule. The larger the number of molecules showing similarity to a given molecule, the less singular the molecule is. Popular molecules within a wide range of singularities are highlighted. For example, raloxifen (**1**), pyrimethamine (**2**) and vemurafenib (**3**) have data in many CC spaces. Likewise, some molecules are more singular than other for which many analogs exist throughout the CC organization (e.g. lovastatin (**4**)).

We then divided each level (A-E) into five sublevels (1-5) corresponding to distinct types or scopes of the data. In total, the CC contains 25 well-defined categories meant to illustrate the most relevant aspects of small molecule characterization. In particular, we stored the 2D (A1) and 3D (A2) structures of compounds, together with their scaffolds (A3), functional groups (A4) and physicochemistry (A5). We also retrieved therapeutic targets (B1) and drug metabolizing enzymes (B2), and molecules co-crystallized with protein chains (B3). We fetched literature binding data (B4) from major chemogenomics databases, and high-throughput target screening results (B5). Moving to a higher order of biology, we looked for ontological classifications of compounds (C1) and focused on human metabolites in a genome-scale metabolic network (C2). In addition, we kept the pathways (C3), biological processes (C4) and protein-protein interactions (C5) of the previously collected binding data. To capture cell-level information, we gathered differential gene expression profiles (D1) and compound growth-inhibition potencies across cancer cell lines (D2). Similarly, we gathered sensitivity profiles over an array of yeast mutants (chemical genetics) (D3), as well as cell morphology changes (high-content screening) (D4). Additional cell sensitivity data available from the literature were also collected (D5). To organize clinical data, we used the traditional ATC classification of drugs (E1), and also drug indications (E2) and side effects (E3) expressed as disease terms, together with therapeutic/adverse outcomes of molecules other than drugs such as environmental chemicals (E4). Finally, we stored drug-drug interactions known to raise pharmacokinetic and efficacy issues (E5).

Further rationale for the choice of the 25 CC categories can be found in Table 1. Overall, we believe that the CC organization is a good representative of what is known of small molecules in the public domain (Table 2). In the Methods section, we extensively describe the data collection protocol. We adopted well-accepted standards, harmonized chemical entries and filtered bioactivities (Figure S1). For example, in the CC D1 space, we discarded those molecules whose transcriptional response was not noticeable and, similarly, only notorious distortions of cell morphology were kept in D4, excluding innocuous compounds. Likewise, we applied target-class specific potency cutoffs to binding data^16^. At the ‘networks’ level (C), we incorporated ontologies and systems biology datasets that are typically outside the scope of compound databases.

**Table 1.** Rationale for the choice of the 5 CC levels (A-E) and sublevels (1-5). Explanations are given from the perspective of drug discovery.

| Level | Name | Rationale |
| --- | --- | --- |
| **A** | Chemistry | Database search and large-scale prediction tools typically use 2D encodings of compounds (A1). Target prediction algorithms often require 3D representations of the compounds (A2), which usually involves an energy-optimization step. In addition, a convenient way to browse the chemical space is through the inspection of scaffolds (A3), and a means to communicate with synthetic chemists is through structural keys (functional groups) (A4). Finally, physicochemical parameters such as the molecular weight are used to rapidly characterize small molecule entities, together with drug-likeness estimations (A5). |
| **B** | Targets | For a relatively small number of molecules (drugs), targets with pharmacological action are known (B1), and drug metabolizing enzymes, transporters, and carriers (B2) are crucial determinants of drug safety (and efficacy). Another prominent set of small molecules are those that have been co-crystallized with protein chains (B3), as they greatly inform of structure-based molecular design. Beyond these, there is a large corpus of binding affinity measurements (B4) and target functional assays (B5) available from the literature and screening campaigns. |
| **C** | Networks | In the mindset of biologists, molecules are put in context via pathways, ontologies and networks. The biological roles of eminent chemical entities are part of an ontology (C1), some entities being metabolites that can be found in the human metabolic network (C2), where substrates and products of enzymes are linked by reactions. These 'higher-order' annotations correspond to a minority of molecules, though. To push systems-level data to the rest of compounds, one can fetch large-scale compound-protein interaction data (e.g. B4), and the canonical pathways (C3) and biological processes (C4) of the proteins may be kept, correspondingly, as annotations for the compound (guilt-by-association). With finer detail, the neighborhoods of these proteins in protein-protein interaction networks can be inspected as well (C5). |
| **D** | Cells | Cell-based assays are bringing about the largest increase in the assortment of data. The LINCS consortium collects gene expression changes after compound dosage (D1), and pioneering platforms such as the NCI-60 continue to produce sensitivity profiles of cancer cell line panels (D2). Similarly, chemical genetics profiles have been proposed to complement genetic interactions in yeast (D3). A very different type of experiment is high-content phenotypic screening, where morphological changes of cells (D4) are measured with microscopy techniques. Growth and proliferation assays (D5) accumulated in the literature over the years can be added to all of these techniques. |
| **E** | Clinics | Clinical data represent the last (and pursued) level of complexity. Drug molecules have been traditionally classified using a therapy/anatomy-oriented hierarchy (E1), although medical vocabularies may be adopted to better defining drug indications (E2) and linking them to disease genetics studies. Likewise, side effects (E3) can be catalogued by parsing drug package inserts and, with a broader scope, there are resources that mine the literature for beneficial and harmful associations between compounds and diseases, including molecules other than drugs such as environmental chemicals (E4). Finally, acknowledging drug-drug interactions (E5) is crucial for medical prescription and the avoidance of undesired |
|  |  | pharmacokinetics and toxicity. |

**Table 2.** Summary of CC spaces and their sources. A brief description of the type of data in each CC coordinate is given. Only major data sources are listed. For more details, URLs and references, please see the Methods.

| Space | Name | Description | Source(s) |
| --- | --- | --- | --- |
| A1 | 2D fingerprints | Binary representation of the 2D structure of a molecule. The neighborhood of every atom is encoded using circular topology hashing. | RDKit |
| A2 | 3D fingerprints | Similar to A1, the 3D structures of the three best conformers after energy minimization are hashed into a binary representation without the need for structural alignment. | E3FP |
| A3 | Scaffolds | Largest molecular scaffold (usually a ring system) remaining after applying Murcko's pruning rules. In addition, we keep the corresponding framework, i.e. a version of the scaffold where all atoms are carbons and all bonds are single. The scaffold and the framework are encoded with path-based fingerprints, suitable for capturing substructures in similarity searches. | RDKit |
| A4 | Structural keys | 166 functional groups and substructures widely accepted by medicinal chemists (MACCS keys). | RDKit |
| A5 | Physicochemistry | Physicochemical properties such as molecular weight, logP and refractivity. Number of hydrogen-bond donors and acceptors, rings, etc. Drug-likeness measurements e.g. number of structural alerts, Lipinski's rule-of-5 violations or chemical beauty (QED). | RDKit and Silico-IT |
| B1 | Mechanisms of action | Drug targets with known pharmacological action and modes (agonist, antagonist, etc.). | DrugBank and ChEMBL |
| B2 | Metabolic genes | Drug-metabolizing enzymes, transporters and carriers. | DrugBank and ChEMBL |
| B3 | Crystals | Small molecules co-crystallized with protein chains. Data are organized on the basis of the structural families of the protein chains. | PDB and ECOD |
| B4 | Binding | Compound-protein binding data available in major public chemogenomics databases. Data come mainly from academic publications and patents. Binding affinities below a class-specific threshold are favored (kinases ≤ | ChEMBL and BindingDB |
| | | 30 nM, GPCRs ≤ 100 nM, nuclear receptors ≤ 100 nM, ion channels ≤ 10 μM and others ≤ 1 μM), and activities at most one order of magnitude higher are kept (capped at 10 μM). | |
| **B5** | HTS bioassays | Hits from screening campaigns against protein targets (mainly confirmatory functional assays below 10 μM). | PubChem Bioassays (from ChEMBL) |
| **C1** | Small molecule roles | Ontology terms associated with small molecules that have recognized biological roles, such as known drugs, metabolites and other natural products. | ChEBI |
| **C2** | Small molecule pathways | Curated reconstruction of human metabolism, containing metabolites and reactions. Data are represented as a network where nodes are metabolites and edges connect substrates and products of reactions. | Recon |
| **C3** | Signaling pathways | Canonical pathways related to known receptors of compounds (as recorded in B4). Pathways are assigned via a guilt-by-association approach, i.e. a molecule is related to a pathway when at least one of the targets is a member of it. | Reactome |
| **C4** | Biological processes | Similar to C3, biological processes from the gene ontology are associated with compounds via a guilt-by-association approach from B4 data. All parent terms are kept, from the 'leaves' of the ontology to its 'root'. | Gene Ontology |
| **C5** | Interactome | Neighborhoods of B4 targets are collected by inspecting several protein-protein interaction networks. A random-walk algorithm is used to obtain a robust measure of 'proximity' in the network. | STRING, InWeb and Pathway Commons, among others |
| **D1** | Gene expression | Transcriptional response of cell lines upon exposure to small molecules. A reference collection of gene expression profiles is used to map all compound profiles using a two-sided gene set enrichment analysis. | L1000 Connectivity Map (Touchstone reference) |
| **D2** | Cancer cell lines | Small molecule sensitivity data (GI50) of a panel of 60 cancer cell lines. | NCI-60 |
| **D3** | Chemical genetics | Growth inhibition profiles in a panel of ~300 yeast mutants. Data are combined with yeast genetic interaction data, so that compounds can be assimilated to genetic alterations when they have similar profiles. | MOSAIC |
| **D4** | Morphology | Changes in U-2 OS cell morphology measured after compound treatment using a multiplexed-cytological 'cell painting' assay. 812 morphology features are recorded via automated microscopy and image analysis. | LINCS Portal |
| **D5** | Cell bioassays | Small molecule cell bioassays reported in ChEMBL. Mainly, growth and proliferation measurements found in the literature. | ChEMBL |
| **E1** | Therapeutic areas | Anatomical Therapeutic Chemical (ATC) classification codes of drugs. All ATC levels are considered. | DrugBank and KEGG |
| **E2** | Indications | Indications of approved drugs and drugs in clinical trials. A controlled medical vocabulary is used. | DrugBank and ChEMBL |
| **E3** | Side effects | Side effects extracted from drug package inserts via text-mining techniques. | SIDER |
| **E4** | Diseases and toxicology | Manually curated relationships between chemicals and diseases. Chemicals include drug molecules and environmental substances, among others. | CTD |
| **E5** | Drug-drug interactions | Changes in the effect of a drug when is co-administered with a second drug. Data are related to pharmacokinetic issues and/or adverse events. | DrugBank |

The CC contains and catalogues information on nearly 800k bioactive compounds (Figure 1B). Evidently, fewer molecules are available as we advance along the CC levels (from A to E), i.e. chemical information (A) is always available (778,460 molecules), whereas clinical data (E) are scarce (9,165 molecules, including 4,232 drugs). The majority of molecules come from the binding literature (B4) or target-based HTS bioassays (B5) (adding up to 705,685 entries in B), and part of this knowledge is transferred to network levels (C3-5) by virtue of biological ontologies, pathways and protein-protein interactions. On a similar scale, the current throughput of cell-based assay platforms (D1-4) is of about 10-20k molecules.

### Signature-based representation of the data

Inspired by the success of chemical descriptors and fingerprints to represent compound structures^17^, we chose to express bioactivity data in a common vector format. Details on how to obtain vectors (*signatures*) for the 25 CC spaces are given in the Methods. In brief, we treated categorical data as sets of ‘terms’, these being proteins, pathways, ATC codes, bit positions of a chemical fingerprint, etc. We then removed frequent and rare terms, and down-weighted the less informative ones (i.e. promiscuous targets, generic biological processes, etc.). Finally, we applied a dimensionality reduction technique, called latent semantic indexing, to ensure that signature components were orthogonal and sorted by their contribution to explaining the ‘variance’ of the data. An analogous procedure (i.e. robust scaling followed by a principal component analysis) was performed on continuous data. For each CC space, we kept the number of components retaining 90% of the variance (Figure S2A). As a result, we obtained 25 numerical matrices, rows corresponding to molecules and columns composing signatures. We named these CC vectors ‘type I signatures’.

Most type I signatures have a length between 500 and 1,500 components (Figure 1B). Longer signatures denote higher complexity or sparseness of the data. Signatures based on gene expression (D1) are the longest, followed by binding data (B4 and B5) and the fine-grained chemical descriptors (A1 and A2). Conversely, physicochemical (A5) and cancer cell line sensitivity signatures (D2) are the shortest. Of note, morphology (D4) signatures only require 26 components to account for the original 812 features, indicating high interdependency of raw measurements (Figure S2B).

We observed that, in all 25 CC spaces, compounds with similar (short-distance) signatures tend to share the mechanism of action and therapeutic area (Figures 1B and S3A). Indeed, bioactivity signatures often correlate better with known mechanisms of action than the more classical chemical signatures (A1-5). Pairwise similarity measurements reveal clusters of molecules and, reassuringly, molecules in the same cluster share targets and therapeutic areas (Figure S3B). More generally, using a similarity-based correlation analysis (Methods) we certified an inter-connection between CC spaces (Figure 1C and S4). Certain links within the CC were expected by design, such as connections within the chemistry spaces (A1-4), or those between binding data (B4) and functionally related versions of them (C3-5). Other correlations have a straightforward interpretation (e.g. drugs with similar targets (B1) have similar indications (E1-2)), and some reflect recognized research biases. For example, we found stronger links between chemistry and mechanisms of action (B1) and therapeutic indications (E1), compared to spaces representing collateral processes such as metabolic enzyme interactions (B2) and toxicology events (E4). Remarkably, we observed correlations between unbiased (omics) datasets. For instance, we found cell sensitivity profiles (D2) to be linked to many CC levels, including the clinical ones (E).

To balance the numerical complexity across CC spaces, we derived an embedded (128-dimension) version of CC signatures (‘type II signatures’). This was achieved by first building similarity networks based on pairwise distances between type I signatures, and then using a network embedding technique to capture (embed) the vicinity of each node (molecule) in a vector space, so that local similarities and more global network properties are seized (Methods, Figures 2A and S5). Figure 2B displays type II signatures for five representative CC datasets, related to drugs used in various disease areas. Visual inspection of these signatures readily highlights some patterns. For example, there is a specific group of side effects (E3) associated with anti-infective drugs, and ophthalmic drugs have similar mechanisms of action (B1) but varied chemistries (A1). We found that, indeed, CC similarity searches greatly increase the chance of identifying drug properties, compared to chemical similarities alone^18^, partly because individual CC spaces have incomplete drug coverage (Figure S6A), and partly because different types of CC data capture different kinds of similarities between drugs (Figure S6B). For example, several CC spaces simultaneously accounted for the relationship between simvastatin, HMGCR inhibition and myocardial infarction (Figure 2D). In the case of doxorubicin, its capacity to inhibit TOP2A was captured by chemical features, while its association with acute myeloid leukemia was identified using transcriptional signatures (D1). For other drugs, like ondansetron, the association with gastroenterology was more trivially provided by target annotations already present in the CC (B1 and B4), whereas for some drugs (e.g. doxifluridine), the chemical similarity to other well-annotated compounds was enough to correctly uncover main therapeutic properties.

**Figure 2.**
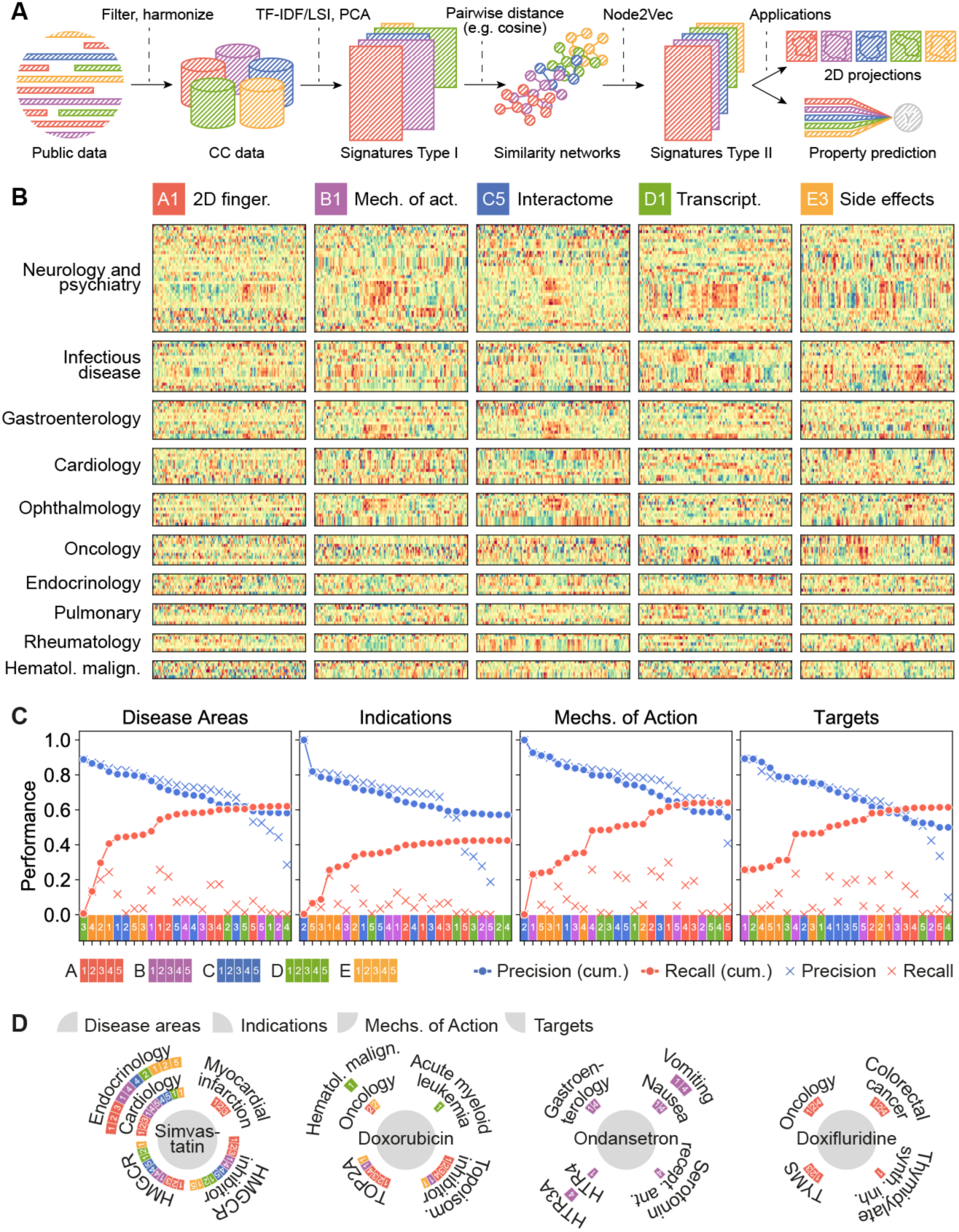
CC signatures visualized. (A) Scheme of the CC pipeline. Public data are filtered, harmonized and unified in the 5×5 CC organization. For each CC space, we obtain type I signatures by doing a (TF-IDF) LSI/PCA dimensionality reduction. With signatures type I, molecules can be compared pairwise to obtain a similarity network. A network embedding algorithm (node2vec) is then applied to derive fixed-length signatures (type II). Type I and/or type II signatures can be used for customary machine learning tasks such as data visualization and property prediction. (B) We plot the numerical values of type II signatures for drugs extracted from the Drug Repurposing Hub^18^, and organize them by disease areas. We chose one illustrative dataset for each CC level, namely A1, B1, C5, D1 and E3. Signatures show, for instance, how chemically unrelated neurological drugs elicit similar patterns of side effects. Likewise, ophthalmological drugs sharing mechanism of action trigger different transcriptional responses. (C) Precision and recall of label predictions (disease areas, indications, mechanisms of action and targets from the Drug Repurposing Hub, Methods). CC spaces are sorted by precision (blue). Recall of molecule-label pairs is shown in red. Dots correspond to cumulative performances (i.e. appending molecule-labels pairs predicted by CC spaces consecutively). Crosses denote individual performances of CC spaces. (D) Examples of true positives, indicating the CC spaces that account for the prediction. Please note that the Drug Repurposing Hub was not included in the CC at the time of compilation.

### Visualizing collections of compounds

CC signatures can be projected to two dimensions (2D), yielding an enhanced characterization of compound libraries (Figure 3A-B). For instance, in Figure 3B we see that, compared to pre-clinical libraries, approved drugs map to a limited area of the physicochemical space (A5), and we can also assess the structural diversity of screening libraries (A4). Beyond chemical structures, we observe that experimental drugs address mechanisms of action (B1) not covered by approved drugs, metabolites or tool compounds. Likewise, we see how they can elicit novel transcriptional changes (D1) and how natural products, such as traditional Chinese medicines (TCMs), may offer new possibilities. We can also notice that a diverse compound collection (Prestwick library) may trigger a limited set of morphological changes (D4). Likewise, in the clinical categories we see the focus of experimental drugs and TCMs (E2), and we also observe differences in the disease landscapes of endogenous and exogenous compounds (E4).

**Figure 3.**
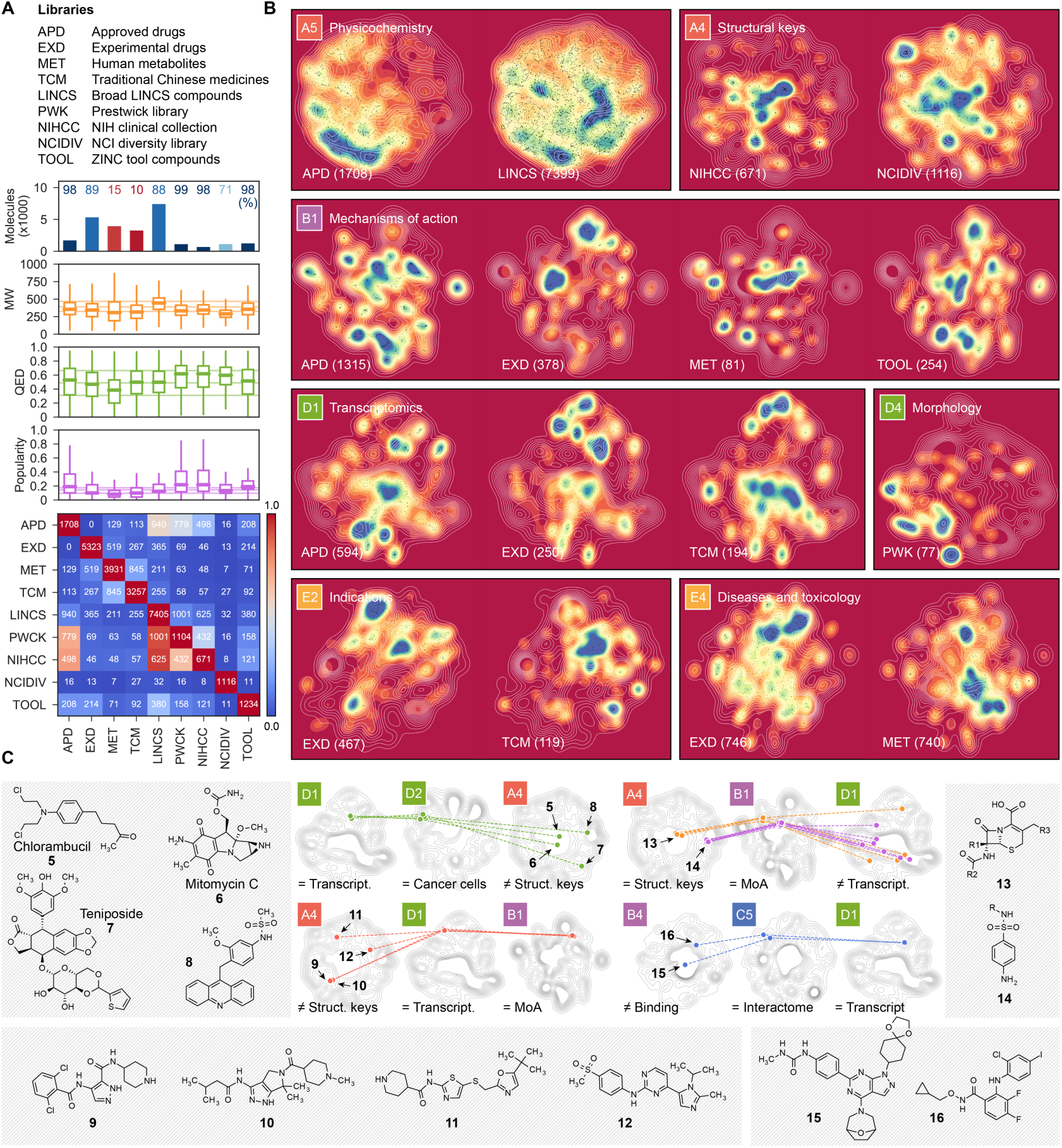
Characterization of compound collections with the CC. (A) Number of CC molecules present in nine representative chemical libraries. The CC coverage of the libraries is shown as a percentage (upper plot), together with molecular weight (MW), chemical beauty (QED)^45^ and popularity scores distributions (middle plot). The overlap between molecules in the compound collections is shown in the heatmap (bottom plot). The libraries contain between 671 (NIHCC) and 7,405 (LINCS) compounds with modest overlap among them. Molecules in the different collections have molecular weights in the 250-500 Da range and comparable indexes of drug-likeness. Virtually all molecules in the APD, PWCK, NIHCC and TOOL collections are catalogued in the CC, while only 10% of the TCM ingredients have a reported bioactivity. As expected, APD are the most popular compounds, followed by the well-annotated chemicals in the PWCK and NIHCC libraries. (B) 2D projections of CC signatures. White lines represent the background distribution of all the molecules in each CC space, and colormaps display the densities of molecules in the indicated collection; in parenthesis, the number of molecules of the collection in the corresponding CC space is shown. (C) Illustrative complex CC queries. Molecules are mapped in more than one CC space, being similar (=) in some of them or different in others (≠). Structures of the selected examples are given.

Further, combining 2D plots throughout the CC facilitates a better understanding of subgroups of compounds, and may inspire complex queries to identify molecules that fulfill multiple characteristics (Figure 3C). For instance, despite being structurally diverse (A4), antitumoral compounds chlorambucil (**5**), mitomycin C (**6**) and teniposide (**7**) trigger similar transcriptional responses (D1) and show similar cell sensitivity profiles (D2), consistent with their known capacity to induce DNA damage—an uncharacterized compound (**8**) was found in this subgroup. We also identified a group of broad-spectrum CDK inhibitors (**9**, **10**, **11** and **12**) that induce a precise transcriptional response (D1). Conversely, we noticed that compounds within antibiotic classes (e.g. beta-lactams **13** or sulfonamides **14**) may be transcriptionally diverse in human cells (D1). Finally, we found compounds (**15**, **16**) targeting kinases (B4) in various signaling pathways (mTOR/PIK3CA and Raf1/MAP2K1/MAP2K2, respectively) that are close in the interactome space (C5) and induce similar cell responses (D1), in agreement with a reported pathway cross-talk with potential for combination therapies^19^.

### Reversion of Alzheimer’s disease signatures

A unique feature of CC signatures is that they can be matched to disease and genetic omics data. For instance, comparison of gene expression signatures in cells can reveal compounds that ‘revert’ transcriptional disease signatures^20, 21^. Typically, these studies require intensive pre-processing^22^, since direct comparisons of gene expression profiles, even within replicas, show modest correlations, and cell-specific biases can confound the analyses^21^. The CC pipeline handles the issues related to multiple doses, time points and cell-lines and returns only one D1 signature per compound (Figure S1).

It is known that the capacity of drugs to revert cancer gene expression profiles correlates with their efficacy^23^ and, indeed, using D1 signatures we obtained similar results on the GDSC panel of cancer cell lines^24^ (Figure S7). The CC spaces are enriched in data obtained from tumoral cell lines and, to evaluate the power of D1 signatures outside the realm of cancer, we engineered novel cells for which no perturbation experiments are available. To this end, we developed cellular models of Alzheimer’s disease (AD) by introducing familial AD (fAD) mutations into SH-SY5Y cells, which are known to recapitulate phenotypes related to neurodegenerative disorders^25^. Using CRISPR/Cas9-induced homology-directed repair, we obtained clones harboring the fAD PSEN1^M146V^ or the APP^V717F^ mutations (Methods and Figures S8A-B). As expected, engineered cells showed an increased extracellular ratio of amyloid β (Aβ) 42 to Aβ40 (Figure S8C), which is a hallmark of fAD mutations^26^.

We measured the transcriptional signatures of PSEN1^M146V^-vs-WT and APP^V717F^-vs-WT cells, which we flipped (reverted) and converted to the CC format (Figure 4A, Methods). Then, we simply ran a similarity search between the CC signatures of the compounds available in D1 and the reverted AD-specific signatures (Data S1). We identified 35 chemically diverse compounds that might have the potential to cancel out transcriptional traits of fAD mutations (Figure S9; Data S1). Of these, three, namely noscapine (**17**) (for the reversion of APP^V717F^ signature), palbociclib (**18**) (for the reversion of PSEN1^M146V^ signature) and the epidermal growth factor receptor (EGFR) inhibitor AG-494 (**19**) (for the reversion APP^V717F^ signature), showed an effect on the secretion of Aβ40 and Aβ42 in SH-SY5Y cells (Figure S8D).

**Figure 4.**
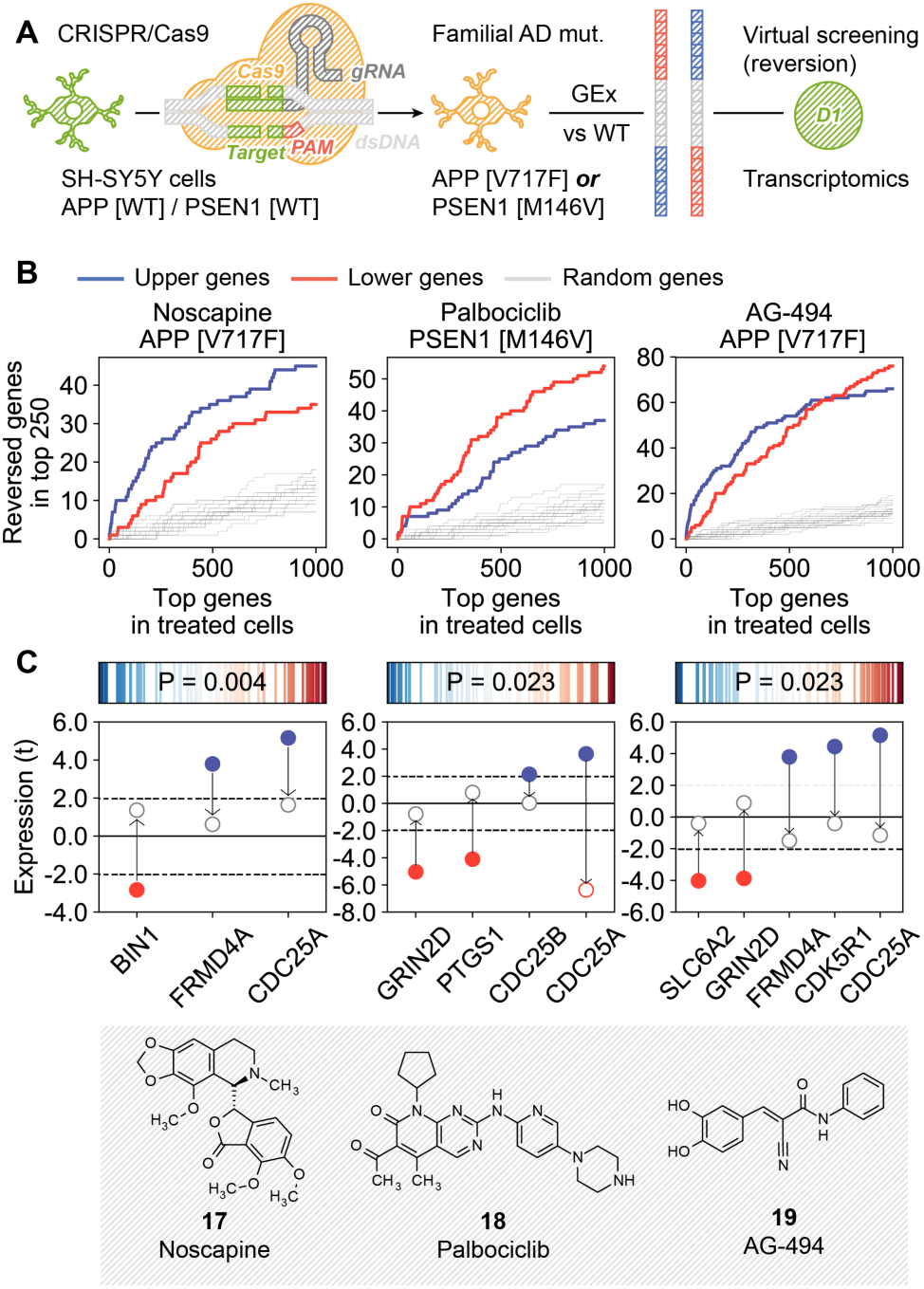
Signature reversion of AD-specific transcriptional profiles. (A) Scheme of the methodology. SH-SY5Y cells were modified with CRISPR to harbor fAD mutations. AD-specific transcriptional signatures were obtained by differential gene expression analysis of mutated-vs-WT gene expression profiles. These signatures were flipped (reversed) and converted to the D1 CC format. Drug candidates were selected based on D1 similarities to the signatures. (B) Experimental results for the three tested candidates, namely noscapine (**17**), palbociclib (**18**) and AG-494 (**19**). In the x-axis, genes are ranked by differential gene expression of treated-vs-untreated mutated cells (APP^V7171F^ or PSEN1^M146V^); this axis relates to both tails of the ranked list (up/down). Correspondingly, in the y-axis we count the number of genes in the mutated-vs-WT signatures that were reverted upon treatment (top 250 genes, up-(blue) and down-(red) regulations). For example, ∼20 of the up-regulated (blue) genes in PSEN1^M146V^ cells are in the top-500 down-regulated genes after treatment with palbociclib, and ∼40 of the down-regulated (red) genes in the PSEN1^M146V^-vs-WT comparison are among the top-500 up-regulated genes when these mutated cells are treated with palbociclib. (C) Reversion of AD-related genes. The upper plots show the tendency of AD genes (according to OpenTargets) to have extreme reversion scores. Reversion scores measure the ratio between ranks in the mutated-vs-WT signatures and flipped (reversed) ranks upon treatment of the mutated cells with the drug. Blue (left of the axis) denotes genes that were up-regulated in the mutated-vs-WT signature and down-regulated upon treatment, and red (right of the axis) denotes genes that were down-regulated in mutated cells and up-regulated upon treatment. The P-value is calculated with a weighted Kolmogorov-Smirnov test based on the absolute value of these reversion scores, i.e. it measures the ‘extremity’ of AD genes. In the bottom plots, we focus on AD genes that were up-(blue) and down-(red) regulated (t-score) in the mutated-vs-WT comparison (bold dots), and we show their expression in the treated-vs-WT comparison (empty dots).

We confirmed that genes up-regulated in SH-SY5Y fAD mutants were indeed down-regulated upon treatment with the drugs, and vice versa (Figure 4B). Moreover, the three drug treatments significantly reverted a subset of genes strongly linked to AD^27^ (Figure 4C), including the recovery of the expression levels of GRIN2D, a glutamate receptor involved in synaptic transmission^28^ and BIN1, a gene involved in synaptic vesicle endocytosis and strongly associated with AD risk^29^.

### Mimicking the activity of biodrugs against IL2R, IL-12 and EGFR

Biodrugs are a family of medicines that includes antibodies and recombinant proteins. Although expensive and prone to having pharmacokinetic issues^30^, biodrugs have the advantage that they bind with high specificity to their targets, which may be proteins considered to be undruggable by small molecule means. We devised a strategy that exploits the signature matching capacity of the CC and identifies compounds that could ‘mimic’ the gene expression profile induced by certain biodrugs (D1), possibly via alternative targets participating in related biological processes (C3-5). After an exploratory analysis (Data S2 and Methods), and based on shRNA interference (knock-down) experiments^21^, we selected three biodrug targets, namely the interleukin (IL)-2 receptor (IL2R), IL-12 and EGFR (Figure 5A).

**Figure 5.**
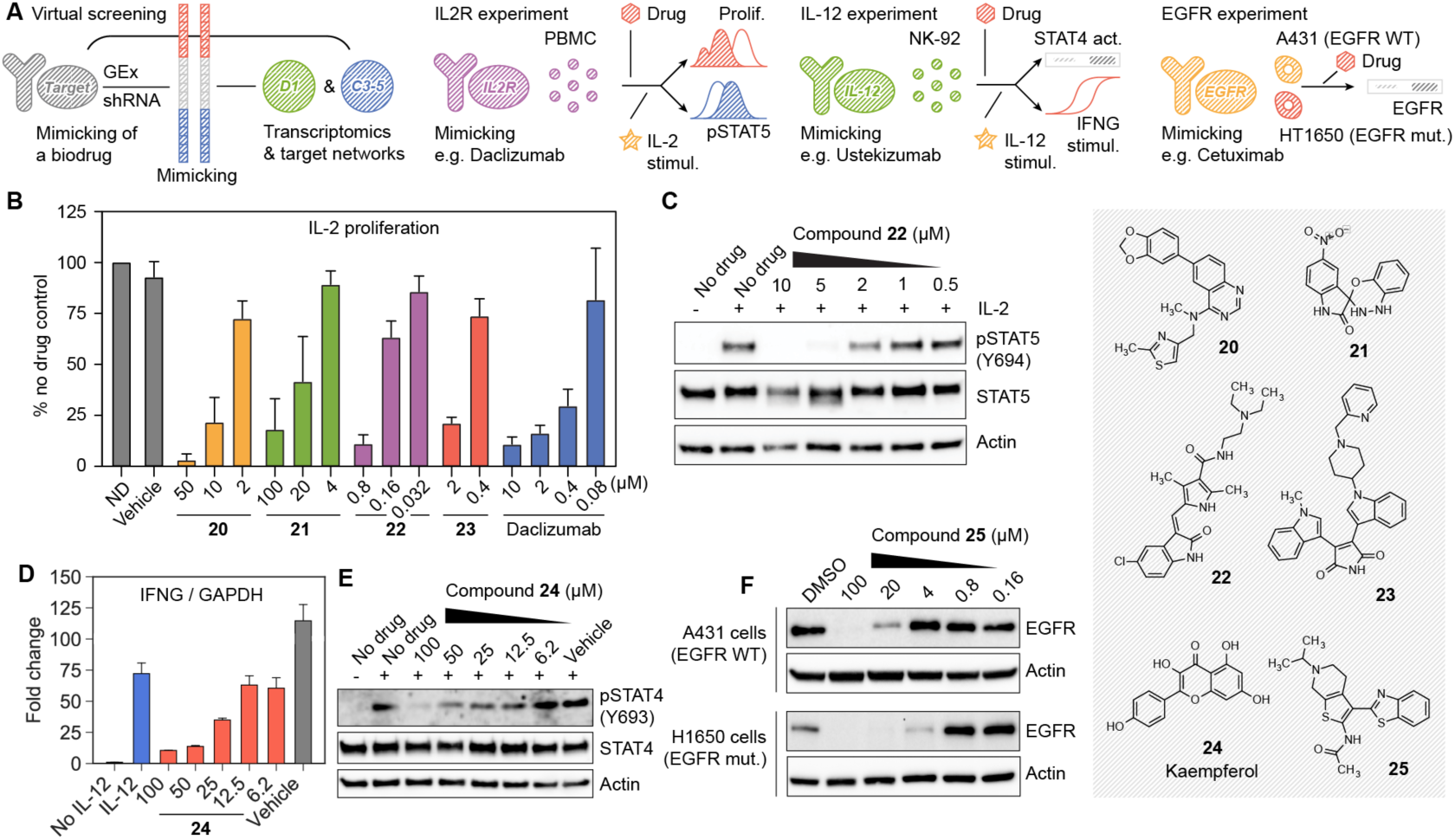
Discovery of chemical analogues of biodrugs. (A) Scheme of the methodology. We look for compounds whose gene expression signatures (D1) would mimic gene expression signatures corresponding to the shRNA knock-down of the target of interest. In addition, we do a networks-level (C3-5) signature matching of the target profiles with those of the compounds. Candidates for IL-2 receptor, IL-12 and EGF receptor are tested in different experimental setups. (B) CD3/CD28 pre-stimulated PBMC were left without treatment for 3 days, labelled with CFSE and then stimulated with IL-2 (0.5 ng/mL) in the presence of the indicated compounds. Three days after stimulation, proliferation was measured by flow cytometry as CFSE label decay and normalized compared to the cells stimulated in the absence of drug (ND). Mean ± SD of 3-5 independent experiments are shown. (C) IL-2-induced STAT5 phosphorylation in PBMC quantified western blot for compound **22**. (D) NK-92 cells were stimulated with IL-12 (50 ng/mL) in the presence of the indicated concentration of compound **24** (kaempferol). *IFNG* mRNA levels after 6 hours were quantified by RT-PCR. Mean ± SD of 3 independent experiments are shown. (E) Phosphorylation of STAT4 at tyrosine 693 was assessed by western blot 1 h after stimulation with IL-12. Total STAT4 and actin antibodies were used as controls. One representative experiment is shown. (F) A431 and H1650 cells were treated for 24 hours with the indicated concentrations of compound **25** (APE1 inhibitor III). We quantified EGFR protein by western blot. Actin was used as a loading control. Representative blots out of three independent experiments are shown.

Daclizumab is a monoclonal antibody targeting the alpha subunit of IL2R, and it is approved for the prevention of organ rejection in transplants. Our computational search highlighted 23 diverse compounds that could potentially mimic the physiological function of daclizumab (Figure S10 and Data S2). We could purchase 19 of these compounds, and we tested their effect in the proliferation of primary human peripheral blood mononuclear cells (PBMC) stimulated with IL-2^31^. Notably, 14 of the molecules showed a significant inhibition of PBMC proliferation induced by IL-2, without substantial effects on cell viability (Figure S11); 13 of the 14 also significantly inhibited PHA-stimulated proliferation^31^ (Table S1 and Figure S12). Note that the hit rate of comparable high-throughput assays ranges between 0.5% and 15% (PubChem BioAssays AIDs: 371, 463, 575, 598, 648, 719, 772 and 2303). When tested in (partially) IL-2 independent cells, the anti-proliferative effect was only moderate (Figure S13). Figure 5B shows confirmatory dose-response curves for four of the candidates, including previously uncharacterized compounds (**20** and **21**).

Further analysis revealed that compound **22** inhibited STAT5 phosphorylation upon IL-2 stimulation (Figure 5C) indicating it is acting in the same signaling pathway blocked by daclizumab. On the contrary, compounds **20**, **21** and **23** did not block STAT5 phosphorylation, suggesting that their anti-proliferative effect is exerted through the blocking of a complementary pathway (Figure S14).

We then sought small molecules that could mimic the function of ustekinumab, a monoclonal antibody whose target is the p40 subunit present in both IL-12 and IL-23 interleukins. Ustekinumab is approved for the treatment of psoriasis, and has potential to ameliorate autoimmune syndromes^32^. One of the best-studied roles of IL-12 is the activation of natural killer (NK) cells, which leads to interferon-gamma (IFNG) production following IL-12 receptor activation and STAT4 phosphorylation. Ustekinumab has been shown to block IL-12-and IL-23-induced IFNG secretion^33^. Our search for compounds that can match C3-5 and D1 signatures of ustekinumab highlighted 17 candidates (Data S2 and Figure S10). We tested the capacity of 11 of them to block IL-12 induced IFNG production in NK cells. One of the compounds, kaempferol (**24**), was indeed able to inhibit *IFNG* transcription in a dose-dependent manner (Figure 5D). Moreover, kaempferol inhibited the phosphorylation of STAT4 in tyrosine 693 in response to IL-12, thereby indicating that this compound exerts its action in an early step of IL-12 signaling (Figure 5E).

Finally, we aimed to mimic the role of monoclonal antibodies that target EGFR for the treatment of advanced colon and head and neck cancers (e.g. cetuximab)^34^. Our CC signature matching search highlighted three candidate compounds (Data S2). Among them, we found apigenin and tanespimycin (17-AAG), which are known to affect EGFR signaling *in vitro* and *in vivo*, and to synergize with cetuximab^35–37^. A third compound was an apurinic/apyrimidinic endodeoxyribonuclease (APE1) inhibitor (**25**) that, to the best of our knowledge, has no reported connection to EGFR. We observed that, indeed, treatment with an increasing dosage of **25** degraded EGFR in wild-type and EGFR^ΔE746-A750^ mutated cells (Figure 5F).

### Similarity searches in the Chemical Checker

We built a web-based resource (CCweb; https://chemicalchecker.org) to facilitate access to our data. As shown in Figure 6, the CCweb displays the 2D projection of each dataset, offering the possibility to use chemical libraries as landmark points and highlighting how individual compounds are distributed and related to each other. In addition, it provides ‘popularity’ and ‘singularity’ (Figure 1D) scores for all compounds, which account for the number of CC spaces related to a certain molecule and the uniqueness (dissimilarity) of a molecule with respect to the rest of compounds, respectively. Moreover, given a molecule of interest, the CCweb retrieves similar molecules in all 25 CC spaces. Most small molecule search engines available to the community are based on chemical similarities, and CCweb is the first to broadly offer search capacity based on biological similarities. CC signatures can be downloaded from the CCweb or simply accessed via a REST API. The entire CCweb resource, including the underlying data and signatures, will be updated every six months.

**Figure 6.**
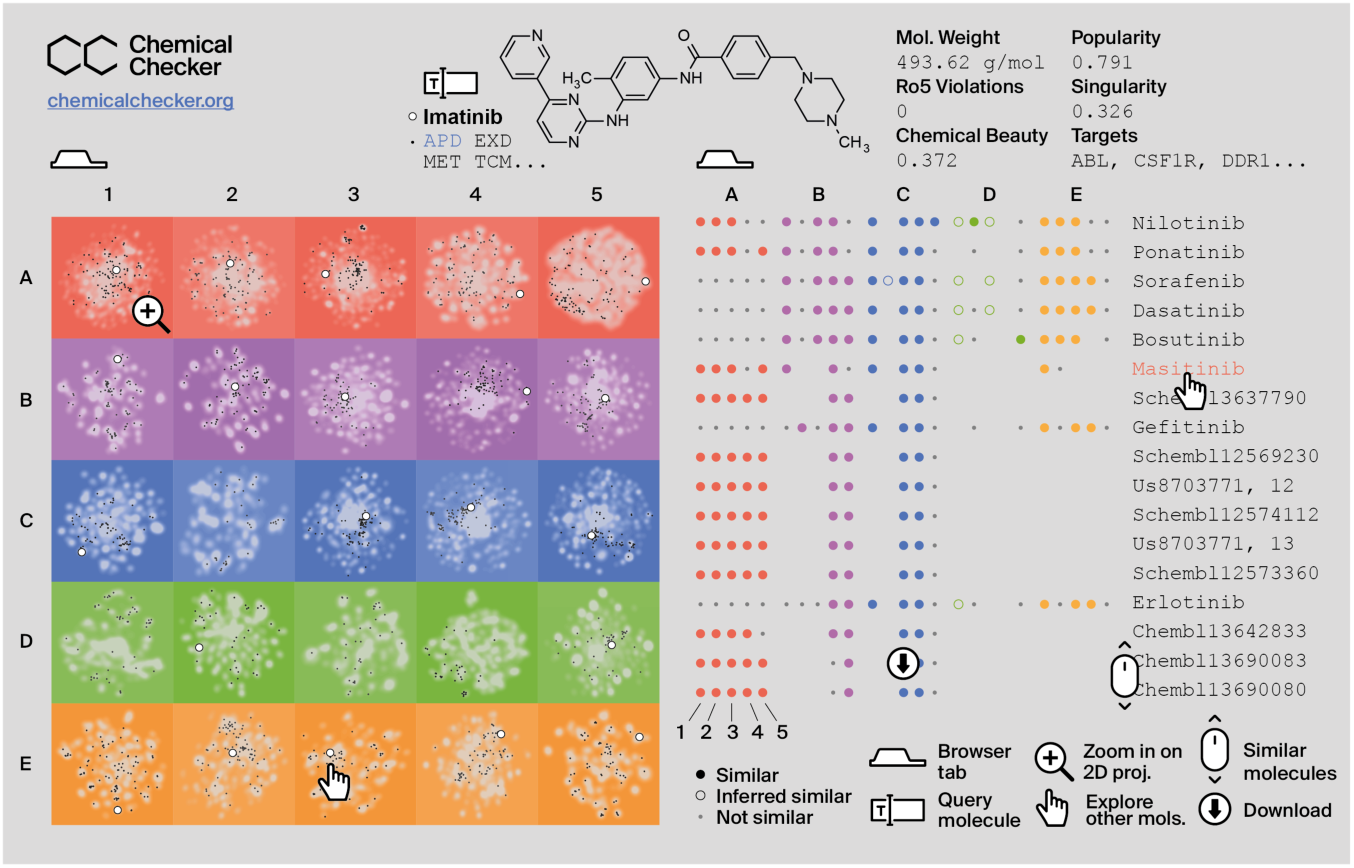
Toy representation of the CCweb resource. The left tab (home page) is an interactive panel of 2D projections, where the query molecule (e.g. imatinib, white dot) can be compared to the CC background (in gray) and to other molecules of interest such as approved drugs (APD, in black). The right tab (exploration page) displays molecules that are similar to the query one. Similarities are measured across the 25 CC spaces (A1-E5).

## Discussion

As small molecule bioactivity data continue to grow in size and diversity, it is essential to present them in a format that is accessible to the majority of researchers. Big initiatives such as OpenPHACTS^38^ and Illuminating the Druggable Genome (IDG)^39^ are undertaking this task, storing links between compounds, genes and diseases in a relational scheme that is ideal for browsing and formulating mechanistic hypotheses. With the CC, we propose an alternative framework based on chemical and biological signatures of compounds. CC signatures are numeric vectors that embed information of a given type (e.g. binding experiments, cell sensitivity profiles or drug side effects) and are suitable for similarity measurements, clustering, visualization and prediction tasks. Such capabilities, we believe, are essential to bridge the gap between relational databases and frontline machine learning algorithms that are able to handle millions of samples but require input data to be expressed in vector format.

The signature-based representation of compounds pushes the similarity principle beyond chemical properties, reaching various ambits of biology. For instance, our preliminary experiments identified candidates to revert AD transcriptional signatures, and we devised a strategy to propose small molecule mimetics of biodrugs. We also used signatures based on pathways, biological processes and networks to gain confidence in our predictions. More generally, we have mapped compound collections to different types of bioactivity spaces, and have shown that similarity searches inside the CC recapitulate drug indications and mechanisms of action.

The current version of the CC contains about 800k molecules. All of them have experimental annotations in at least one of the biological levels (B-E, typically B4 and B5). The known chemical space is much larger than this, containing millions of commercial compounds and a cosmic number of synthetically accessible virtual molecules^40^. A good proportion of the molecules will not be bioactive, falling outside the scope of the CC. However, it is recognized that the bioactive chemical space remains mostly uncharted^41^. Given the remarkable correlations observed within the CC, we are confident that our resource will be coupled to signature inference in the future, providing a means to rapidly characterize any molecule of interest.

## Online Methods

### Raw data

Small molecule entries were collected from several resources (Table 1 and Figure S1) and stored by standard InChIKey. The InChIKey is a 25-character string that encodes the connectivity of the molecule (first 14 characters), other details like stereochemistry (next 8 characters), the kind and version of the key (next 2 characters), and the protonation state (last character). To assign an InChIKey to each small molecule, we read the structure as given by the source database (usually a SMILES string) and followed a standardization procedure consisting of salt and solvent removal, charge neutralization and the application of rules to tautomeric groups (https://github.com/flatkinson/standardiser).

#### A: Chemistry

##### A1: 2D fingerprints

2048-bit Morgan fingerprints (radius = 2) were calculated using the RDKIT (http://rdkit.org).

##### A2: 3D fingerprints

1024-bit E3FP fingerprints (https://github.com/keiserlab/e3fp) were calculated by merging the results of the three best conformers obtained with a UFF energy minimization, as recommended in the E3FP publication^42^.

##### A3: Scaffolds

We extracted the Murcko’s scaffold of each molecule^43^. In addition, we derived the molecular framework of the scaffold, i.e. all heavy atoms were converted to carbon atoms and all bonds were simplified to single bonds. When no scaffold could be obtained, we kept the full structure of the molecule and the corresponding framework. 1024-bit Morgan fingerprints (radius = 2) were then calculated for each molecule and concatenated in a 2048-bit fingerprint.

##### A4: Structural keys

The widely used, human-readable MACCS 166-keys^44^ were calculated using the RDKIT. MACCS keys represent structural features relevant to medicinal chemistry. Each key is associated with a SMARTS pattern.

##### A5: Physicochemistry

For each molecule, we calculated the molecular weight, number of heavy atoms, number of heteroatoms, number of rings, number of aliphatic rings, number of aromatic rings, logP, molecular refractivity, number of hydrogen bond (HB) acceptors, number of HB donors, polar surface area and number of rotatable bonds. In addition, we flagged the structural alerts proposed by Hopkins and coworkers^45^ and those listed in ChEMBL^4^ (v22, https://www.ebi.ac.uk/chembl). We also counted Lipinski’s rule-of-5 violations^46^ and rule-of-3 violations^47^. Finally, the chemical beauty (QED)^45^ was quantified using the Silicos-IT kit (http://www.silicos-it.com).

#### B: Targets

##### B1: Mechanism of action

Mechanisms of action of approved and experimental drugs were collected from DrugBank^48^ (v4, https://www.drugbank.ca) by selecting those protein targets with a known pharmacological action and action mode. Similarly, we fetched from ChEMBL those drugs with a known mode of action. We distinguished between ‘activation’ modes (agonist, activator, etc.) and ‘inhibition’ modes (antagonist, competitor, etc.). Together with the identity of the protein targets, we retained protein class memberships (GPCRs, kinases, etc.) at all levels of the ChEMBL target hierarchy.

##### B2: Metabolic genes

We collected drug-metabolizing enzymes, transporters and carriers from DrugBank. To these, we added proteins involved in drug metabolism as recorded in ChEMBL. As in B1, we retained protein class information.

##### B3: Crystals

We downloaded ligand data from the Protein Data Bank (https://www3.rcsb.org, February 2017). Protein structures bound to each small molecule were then annotated with family (F- and T-groups) and superfamily (H- and X-groups) information, following the Evolutionary Classification of Protein Domains^49^ (ECOD v1.4, http://prodata.swmed.edu/ecod).

##### B4: Binding

Protein binding data were obtained from ChEMBL by searching for bioassays of ‘binding’ type, related to ‘single proteins’ with an experimental measure of standard type (‘pChEMBL’ value available). We also collected BindingDB records with activity expressed as concentrations^50^ (https://www.bindingdb.org, February 2017). Data were discretized by applying the following activity cutoffs, recommended in Pharos^16^ (http://pharos.nih.gov/idg): kinases ≤ 30 nM, GPCRs ≤ 100 nM, nuclear receptors ≤ 100 nM, ion channels ≤ 10 µM and others ≤ 1 µM. We also kept activities one order of magnitude lower than the class-specific cutoff (to a maximum of 10 µM), and gave these annotations half the weight in downstream analyses (i.e. log10 scaling). Finally, protein class hierarchy information was kept as in B1.

##### B5: HTS bioassays

The largest public repository of small molecule screening data is PubChem Bioassays^5^. Bioactivity values from this repository were directly downloaded from ChEMBL, since the latter conveniently applies a processing pipeline that collects only confirmatory assays and maps related protein targets to UniProt identifiers. Most of the assays belong to the ‘functional’ category. For completeness, we included other functional assays available in ChEMBL. We chose a relaxed activity cutoff of 10 µM, or checked for the word ‘active’ in the description of the assay. We kept the protein class hierarchy as in B1.

#### C: Networks

##### C1: Small molecule roles

We downloaded the Chemical Entities of Biological Interest (ChEBI) ontology^51^ (v150, http://www.ebi.ac.uk/chebi). Only ‘3-star’ molecules were considered. The ‘role’ ontology was loaded as a directed graph capturing ‘is a’, ‘is conjugate acid/base of’, ‘is enantiomer of’, ‘is tautomer of’ and ‘has role’ relationships. In the ChEBI graph, molecules are ‘leaves’. We searched for paths to reach the ‘root’ of the graph (i.e. the ‘role’ node) from each of the leaves. Terms belonging to these paths were annotated to the corresponding molecules.

##### C2: Metabolic pathways

We downloaded the reconstruction of human metabolism (Recon)^52^ from Pathway Commons^53^ (http://www.pathwaycommons.org, July 2017) in binary interaction form. Data were represented as an undirected graph where nodes are metabolites and edges denote reactions. We then computed an ‘influence matrix’ based on this metabolic network. In brief, positions in the influence matrix quantify the proximity between pairs of metabolites. The neighbors of each molecule were selected following a weighting scheme to favor proximal metabolites. Please see C5 below for more details on how influence matrices are calculated and neighbors extracted and weighted therefrom.

##### C3: Signaling pathways

The C2 space above is focused on endogenous metabolites. Conversely, this C3 space (and the C4 and C5 spaces) is aimed at any molecule with known protein targets. In this case, we list the biological pathways that may be affected by the interaction of a molecule with its targets. Human pathways were collected from Reactome^54^ (https://reactome.org, May 2017), and we chose to use binding activities from B4, since this is an extensive dataset containing mostly literature data with well-accepted activity thresholds. In B4, 24.5% of the compound-protein interactions do not correspond to human proteins. These were mapped to their human orthologs using MetaPhOrs^55^ (http://orthology.phylomedb.org, May 2017), following the observation that binding activities can be safely transferred between orthologous proteins^56^, especially if they belong to closely related species, as it is the case for B4 data^57^. Of all the non-human proteins mapped to the human orthologs, 94.4% were mammal proteins.

Molecules were annotated with Reactome pathways using a simple guilt-by-association approach, i.e. a pathway was kept when at least one of its proteins was a target of the molecule. Pathways at all levels of the Reactome hierarchy were evaluated, and weight was given to the pathway annotation on the basis of the compound-target binding record (see B4).

##### C4: Biological processes

. We downloaded the Gene Ontology Annotation (GOA) database (https://www.ebi.ac.uk/GOA, May 2017) and read the ‘biological process’ (BP) branch of the ontology as a directed acyclic graph (DAG) (‘is a’ relationships). Proteins were annotated with their GOA BP terms plus parent terms (up to the root of the DAG). Similar to C3, we associated molecules with BP terms by simply checking the annotations of the molecule targets (B4).

##### C5: Interactomes

We collected five representative protein-protein interaction (PPI) networks, namely STRING (score > 700, i.e. high confidence)^58^ (v10, https://string-db.org) [14,725 proteins (*p*), 300,686 interactions (*i*)], InWeb (score ≥ 0.5)^59^ (http://www.intomics.com/inbio/map, March 2017) [10,100 *p*, 168,970 *i*], a portion of Pathway Commons containing interactions from known pathways (KEGG^60^, NetPath^61^, PANTHER^62^ and WikiPathways^63^) [9,344 *p*, 242,962 *i*], an in-house network of physical binary PPIs^64^ [13,038 *p*, 64,659 *i*], and a network of metabolic genes based on Recon (v2, http://vmh.uni.lu) [1,628 *p*, 246,937 *i*]. To build this last network, we linked two metabolic proteins (enzymes or transporters) when the product metabolite of the first was the substrate of the second, or when both were needed to perform a certain reaction, suggesting that they are part of the same protein complex. Edges between proteins were weighted inversely proportional to the number of reactions involving their shared metabolites, so that ‘currency’ metabolites such as ATP and water had marginal impact on the network connectivity. In order to control for indirect associations, we deconvoluted the network^52^ using edge weights and setting a network deconvolution score cutoff of 0.9.

The five networks above were treated separately in the following procedures. Given a PPI network containing *n* nodes, we computed a *n* x *n* ‘influence matrix’ using HotNet^65^ (v2, https://github.com/raphael-group/hotnet2). The influence matrix measures how likely a random walker departing from node *©* is to reach node *j*. This measure accounts for topological features such as centrality and betweenness, hence it is a more robust quantification of the relationship between nodes than the simple presence or absence of an interaction between them. Then, given the targets of a small molecule (B4), we looked in the matrix for the nodes that are most ‘influenced’ by these targets, i.e. we retrieved proteins other than the target that are likely to be affected by the compound. The search for ‘influenced’ nodes was done as follows. First, non-diagonal values in the influence matrix were scaled from 0 to 10 and expressed as integers; as expected, most of the values were equal to 0, meaning that most proteins pairs were not influencing each other. Then, for each target of a certain compound, we kept proteins with a non-0 influence score (the target itself was given a score of 10 and, when one protein was influenced by more than one target, the maximum score was kept). Finally, these scores were multiplied by the weight of the compound-target annotation (see B4). As a result, for each small molecule in each network, we obtained a weighted set of proteins that may be affected by the interplay with the targets. Results from the five different networks were concatenated for further analyses.

#### D: Cells

##### D1: Gene expression

Transcriptional profiles of treated cultured cells were obtained from the L1000 Connectivity Map^21^ (Phase I: GSE92742 and Phase II: GSE70138 in the Gene Expression Omnibus, March 2017). In this dataset, each ‘perturbagen’ (small molecule, shRNA or overexpressed gene) has several gene expression signatures assigned, corresponding to different doses, times of exposure, cell lines, etc. We took level 5 (replica-aggregated) signatures, considering both landmark and inferred gene expressions. Signatures with a low correlation between replicas (‘distil_cc_q75’ < 0.2) were discarded. Following the authors’ recommendations^21^, we picked an ‘exemplar’ signature for each perturbagen in each available cell line by prioritizing signatures with a number of samples between 2 and 6, and selecting the one with a highest transcriptional activity score (TAS). As a result, each perturbagen-cell line pair has one (and only one) signature assigned.

After the filtering above, the complete L1000 Connectivity Map contained 22,118 perturbagens, each of them tested, on average, in 3.8 of 86 cell lines. A smaller, functionally diverse, and well-annotated subset of the data is the Touchstone dataset, which is focused on 8,880 perturbagens screened against a core collection of 9 cell lines. The Touchstone dataset is the one that is queried in the online application of the L1000 Connectivity Map (https://clue.io/l1000-query), and we chose to use it as a reference collection of signatures. Accordingly, we measured pairwise similarities (‘connectivities’) between the small molecule signatures (‘trt_cp’) of the full dataset (F) and the Touchstone (T) signatures (‘trt_cp’, ‘trt_sh.cgs’ and ‘trt_oe’). To this end, we took the top 250 over- and under-expressed genes of the F-signature^66^ and ran a two-way gene-set enrichment analysis (GSEA) against T-signatures to obtain connectivity scores (CS)^21^ corresponding to the average between the GSEA enrichment score (ES) of up-regulated genes and the GSEA of down-regulated ones.

Connectivity scores were then normalized (NCS) so that they were comparable between T-signature cell lines and perturbation types (small molecule, shRNA or gene over-expression). Normalization was simply done by dividing CS by its average in each perturbation type category.

The CC is compound-centric, hence we summarized the results above, obtained for individual cell types, into a single measure of connectivity between F-molecules and T-perturbagens. A cell-summarized (consensus) connectivity score (NCS_cons_) was given by the maximum tertile statistic, first across T-signatures and then across F-signatures.

As a result, we obtained a F-vs-T connectivity matrix comparing the expression patterns of all molecules to the expression patterns of reference (Touchstone) perturbagens. Finally, we discretized this connectivity matrix by selecting, for each F-molecule, significantly similar T-perturbagens (P < 0.01, i.e. 99% percentile of NCS_cons_). Molecules with less than 5 significantly similar T-perturbagens were discarded.

##### D2: Cancer cell lines

Modern cancer cell line panels such as the Cancer Cell Line Encyclopedia (CCLE)^67^, the Genomics of Drug Sensitivity in Cancer (GDSC)^24^ and the Cancer Therapeutics Response Portal (CTRP)^68^ contain about a thousand cell lines but are short on screened molecules, having at most a few hundred of them. Conversely, the more classical NCI-60 cancer panel^69^, while significantly narrower (60 cell lines), has almost 20k molecules screened for sensitivity, thus making it a better case for the CC. We collected z-transformed GI50 data from the NIH Developmental Therapeutics Program (https://dtp.cancer.gov, June 2016). Only molecules screened against at least 50 of the cell lines were considered. When more than one sensitivity profile was available for a given InChIKey, we kept the one with the largest number of assayed cell lines. This left us with a small molecule sensitivity matrix that was 95.2% complete. Missing values were imputed using the MICE imputation algorithm over 100 iterations^70^.

##### D3: Chemical genetics

We downloaded chemical genetics data from MOSAIC^71^ (http://mosaic.cs.umn.edu, September 2017). The raw chemical genetics dataset contains ∼10k small molecules screened against ∼300 yeast mutants. These ∼300 yeast mutants were selected by the authors of the dataset so that they are representative of a broader panel of ∼5k mutants. The ∼10k x ∼300 chemical genetics matrix then becomes truly informative when it is compared to the ∼5k x ∼300 genetic interaction matrix, in such a way that similarities between compounds and gene alterations can be discovered. This comparison is conveniently published in MOSAIC as a ‘gene target prediction’ file. We discretized the information in this file by keeping the identity of yeast mutants whose profiles had a similarity score above 7.12 (corresponding to a P-value of 0.001) and, with half the weight, yeast mutants with a score above 3.37 (P-value of 0.01). Only 3,560 molecules passed this significance filtering.

##### D4: Morphology

We downloaded the LDS-1195 dataset from the LINCS Data Portal (http://lincsportal.ccs.miami.edu), corresponding to cell painting morphological profiles^72^. This dataset reports 812 cell image features measured after treatment of cells with ∼30k compounds. In order to filter out molecules that do not have a substantial impact on cell morphology, we first counted the number of features (Nf) of each molecule that were significantly extreme (P < 0.01, i.e. bottom 1% and top 99% of feature value distribution). We then repeated the same procedure to column-wise permuted versions of the data, and kept the Nf_0_ point of P < 0.01 significance of this null distribution. Accordingly, we considered that molecules with Nf < Nf_0_ did not trigger a significant morphological pattern, and we consequently discarded them; 12,075 molecules remained after the filtering.

##### D5: Cell bioassays

We downloaded literature cell bioassay data from ChEMBL. We kept only standardized activity data given in commonly used units such as GI50, LC50 or IC50. Activities below 1 µM were retained, together with values beyond the 50% when data were percentual. We excluded cell lines that could not be mapped to the Cellosaurus ontology (v22, https://web.expasy.org/cellosaurus). The Cellosaurus was used to identify and retain ‘derived from’ relationships between cell lines.

#### E: Clinics

##### E1: Therapeutic areas

We collected Anatomical Therapeutic Chemical (ATC) classification system codes from DrugBank and KEGG. To capture the ATC hierarchy, we annotated molecules with their full ATC code (level 5), plus all higher levels (4 to 1).

##### E2: Indications

We fetched approved and phase I-IV drug indications from ChEMBL and RepoDB^73^ (v1, http://apps.chiragjpgroup.org/repoDB). RepoDB is an indication-oriented version of DrugBank. UMLS disease terms in RepoDB were mapped to the MeSH vocabulary using DisGeNET^74^ (v4, http://disgenet.org) (MeSH is the preferred vocabulary in ChEMBL). We considered approved drug indications, together with those in clinical trials, from both databases. When a drug was indicated for more than one disease, weight was assigned to each indication depending on the clinical status (phase I to phase IV/approved), so that e.g. phase II annotations were twice as weighted as phase I’s. MeSH terms were spanned across the MeSH hierarchy as explained in E4 below. We kept the maximum weight for each parent term.

##### E3: Side effects

We collected drug side effects from SIDER^75^ (v4, http://sideeffects.embl.de), expressed as UMLS terms. We did not consider frequency information since we and others have found it to be too scarce for comprehensive statistical analyses^76, 77^.

##### E4: Disease phenotypes

Associations between chemicals and disease phenotypes were downloaded from the Comparative Toxicogenomics Database (CTD)^78^ (http://ctdbase.org, July 2016). We took only ‘curated’ CTD data. In CTD, compound-disease associations are classified as ‘therapeutic’ (T) or ‘marker/mechanism’ (M) (usually corresponding to a disease-causing effect). T and M annotations were kept separately for each molecule. CTD contains a medical vocabulary (MEDIC) that is essentially based on the MeSH hierarchy. For each annotated disease, we added parent terms all the way to the root of the MEDIC hierarchy.

##### E5: Drug-drug interactions

To the best of our knowledge, DrugBank is the largest, most reliable drug-drug interaction (DDI) repository^79^. DDI data was directly downloaded directly from this database.

### Type I CC signatures

#### ***Discrete (and discretized) data*** *(A1-4, B1-5, C1-5, D1, D3, D5, E1-5)*

Discrete data are expressed as sets of *terms*, where terms can be proteins, pathways, ATC codes, bit positions of a chemical fingerprint, etc. In some CC spaces, terms are weighted according to their quality or importance (e.g. B4 or C5). In order to convert these sets of terms to a vector form, we applied a protocol originally developed for the numerical representation and comparison of text documents. First, we removed infrequent and frequent terms, i.e. terms occurring in less than 5 and more than *m* of the molecules (*m* = 80% for A1-3, *m* = 90% for A4 and *m* = 25% for the rest). We then applied a TF-IDF transformation to the terms of each molecule, so that ‘term frequency’ was proportional to the weight of the term (when applicable; 1 otherwise), and the ‘document frequency’ corresponded to the occurrence of the term along the corpus of molecules. As a result of the TF-IDF transformation, less informative terms (i.e. promiscuous targets, generic BPs, etc.) become less important. Finally, we applied Latent Semantic Indexing (LSI) to the TF-IDF-transformed corpus. LSI is a dimensionality reduction technique based on singular-value decomposition (SVD), hence it has parallelisms with the more popular principal component analysis (PCA). In particular, LSI components are also orthogonal and sorted by their contribution to explaining the ‘variance’ of the data. For each dataset, we kept the number of LSI components that explains 90% of the variance. The resulting signatures are thus comprehensive. We also kept track of the ‘elbow’ point of the variance-explained curve (i.e. the point of maximum curvature in the scree plot), as this point gives a good trade-off between accuracy (high dimensions) and interpretability (low dimensions).

#### Continuous data (A5, D2, D4)

Data of this type were first robustly scaled column-wise (median = 0, median absolute deviation = 1; capped at +/- 10). Then, for each CC space, we performed a PCA and chose the number of components that explained 90% of the variance. The elbow point was also kept.

### Type II CC signatures

We built 25 similarity networks (nodes: molecules, edges: similarities (empirical – log_10_*P*-value)). We kept only similarities below a significance *P*-value of 0.01, and, for each node, we considered at maximum 100 links to other nodes. We ensured that each node was connected to at least 3 other nodes (ranked by similarity).

We then ran node2vec^80^ to obtain *embeddings* for each node (molecule) in each network. Node2vec was run with default parameters, i.e. p = 1, q = 1, k = 10 (context size), r = 10 (walks per source), l = 80 (length of walk). We found an embedding dimension of 128 to be a robust choice across CC spaces (Figure S5).

### Clustering and 2D projections of CC signatures

#### Clustering

We used a product-quantized (PQ) version of k-means^81^ (using PQ-table lookups, PQ-encoders of 256 bits and 8 vector splits) to cluster molecules based on signature similarities. The k-means algorithm requires that a number of clusters *k* is pre-defined. We ran k-means with *k* in the range 2 < *k* < *√N*, *N* being the number of molecules in the CC space. Inertia (sum of sample distances to centroids) and concentration (inverse of dispersion; i.e. number of centroids with another centroid at a significantly close distance (P-value < 0.05)) were calculated at each *k.* Inertia and concentration curves were smoothed with the Hanning method (window length of *√N*/ 10) and scaled between 0 and 1 within the explored *k* range. We chose a *k* that maximized the geometric mean of both curves, weighting the dispersion curve by the length of the signature with respect to *√N*/ *2*.

#### 2D projections

In order to have 2D projections of comparable granularity across CC spaces, we performed a k-means clustering on spaces with more than 1,000 samples and took a *k* of N/2, capped at 15,000. Then, projections shown in the CCweb and figures of the paper were performed with type I signatures (we observed very similar results with type II signatures). Signatures were projected in a 2D plane using the Barnes-Hut t-SNE algorithm^82^ with a perplexity of 30 and an angle of 0.5. HDBscan^83^ was used to identify sparse points (outliers) in the projection. After removing these points, t-SNE was re-run. When necessary, samples were assigned the coordinates of their corresponding centroid.

### Correlation between CC spaces

To measure the correlation between two CC spaces, we checked whether, according to the respective CC signatures, molecules in the first space are also similar in the second. We designed a composite correlation coefficient (κ) that quantifies the agreement between the two ranked-similarity lists in several ways (Figure S4). The κ coefficient includes a canonical correlation analysis as well, based on the analysis of dataset cross-covariance and the identification of maximally-correlated linear combinations of the two signatures. Thus, high κ values indicate that two CC datasets share similar-molecule pairs and that ‘common directions’ can be found between signatures of two datasets.

#### Canonical correlation analysis

. Given two CC spaces *X* and *Y* (paired rows, with *m x p* and *m x q* dimensions, respectively, where *m* is the number of common molecules in both CC spaces and *p* and *q* are their signature lengths), we did a canonical correlation analysis (CCA) to identify canonical variables (i.e. linear combinations of signature components) that optimally correlate. Correlation between datasets was measured by averaging the Pearson’s correlation between the two first components identified.

#### Rank-biased overlap

We measured the rank-biased overlap (RBO)^84^ between two sorted similarity lists. RBO simulates the behavior of a user scrolling down a list of search results in the web. Higher probabilities of ‘visiting’ a search result are given to higher similarity scores. Two CC spaces with similar RBO lists are thus correlated.

#### Discretized similarity

When comparing CC spaces pairwise, we classified similarities in the P-value intervals < 1e-5 (i.e. ∼0), 0.001, 0.01, 0.1, 0.25 and > 0.25. Pairs at these intervals were counted in an ordinal contingency table. Counts were L1-normalized row-wise and column-wise iteratively, and a kappa correlation score was measured on the contingency table using the standard quadratic weighting.

#### Cumulative conditional probabilities

. Likewise, we calculated conditional probabilities of two molecules being similar in one CC space when a similarity is observed in another space. The area under the cumulative conditional probabilities (log2-scaled) can be used as a measure of correlation between two spaces.

#### Consensus measure

. The correlation measures explained above were unified to a single dataset correlation measure (κ) by simply taking the median value of the individual correlation measures. Values of individual correlations were quantile-normalized prior to this computation.

### Label assignment based on similarity searches

We downloaded drug annotation data from the Drug Repurposing Hub^18^ (March 2019). We mapped 5,880 drugs to the CC. These were related to 24 ‘Disease Areas’, 664 ‘Indications’, 1,067 ‘Mechanisms of Action’ and 2,249 ‘Targets’. We then devised the following ‘label assignment’ exercise. For each molecule, we looked for the similar molecules in the dataset using each of the 25 CC spaces separately. Similar molecules were defined as having an empirical P < 0.001, calculated on a background specific to the Drug Repurposing Hub. At least 3 (and at most 10) neighbors (similar molecules) were considered per molecule. Then, we evaluated the enrichment (Fisher’s exact test, P < 0.001) of labels among the neighbors of the molecules^85^. We required a label to be represented at least in 5 molecules in the dataset and, at least, in 3 of the neighbors of the molecule.

For each CC space, we performed independent label assignment exercises for all molecules with known labels. Precision and recall were evaluated, both for CC spaces individually and in a cumulative manner by aggregating (appending) the predictions of each CC space sequentially, spaces being sorted by individual precision.

### Chemical Checker web resource

The CCweb resource (https://chemicalchecker.org) is a tool to explore the bioactivity of small molecules, focusing mainly on the identification of compound similarities. To keep pace with its source data, the CCweb will be updated every six months. A simplified representation of the website can be found in Figure 6.

#### Home page

##### Panels

The main page of the CCweb consists of a 5×5 grid of panels displaying 2D projections of the signatures (A1-E5). The distribution of *all* molecules in each CC space is shown as a gray density plot. The user can click on a panel to amplify it. Small molecule counts and a short explanation of the selected CC space accompany the plot.

##### Query and molecule card

Molecules can be queried by InChIKey, PubChem Compound ID (CID) or name. If found in the CC, the compound is shown in the 2D panels where data are available for it. On the right side of the page, a ‘molecule card’ gives basic information about the compound of interest (molecular weight and formula, rule-of-5 violations, chemical beauty, popularity, singularity, etc.). Of note, a list of targets is given. This list is not meant to be comprehensive, and we encourage users to visit dedicated databases such as ChEMBL to learn more about the targets of their molecules of interest. Targets are sorted by species (human first), then by source (B1 > B4 > B2 > B5) and potency (in the case of B4), and finally in alphabetical order.

##### Libraries and landmark molecules

To facilitate navigation of the 2D panels, we offer the possibility to overlay molecules from the popular chemical collections discussed in this article (approved drugs, Prestwick library, etc.; see Figure 3). These collections can be chosen with the ‘change’ button on the left of the screen. We have selected 100 ‘landmark’ molecules from each collection, since in most cases displaying the full library would be impractical. The selection of the 100 landmark molecules is done such that they are present in as many panels as possible, and favoring their distribution in the 2D projections. To achieve this, we start with the molecule with the highest popularity score. Molecules are sequentially added by bagging always from the most ‘orphan’ CC spaces (i.e. the datasets that have the fewest molecules selected). From these, we consider only molecules from the most ‘orphan’ clusters and, among the remaining candidates, we select the one with the highest popularity (i.e. more spaces available).

In summary, in the home page of the CCweb, the user can query small molecules and obtain an overview of their location inside the CC. The user will learn the CC spaces where these molecules have data available, with gray 2D density plots indicating whether they are ‘peripheral’ (low-density regions) or ‘central’ (high-density regions). To have a better sense of the location of query molecules, landmark compounds from popular collections can be displayed. Deeper insights can be obtained by clicking on the ‘explore’ button for a molecule of choice.

#### Explore page

##### List of similar molecules

When a molecule is ‘explored’, we look for similar molecules in the CC database and display them in a 25-column table, corresponding to the CC spaces. In CC spaces where the molecule is present, we measure similarities to other molecules in the space. If the molecule is not present in the space, we infer similarities only against molecules that are (absent-vs-present). Inference of similarities is done by simple probabilistic rules, based on the©e Bayes formulation; i.e. conditional probabilities of being ‘similar’ (P < 10^-5^, < 0.001, < 0.01, < 0.05, < 0.25, > 0.25) in an observed space, based on ‘observed’ similarities© naive Bayes formula was modified by weights^86^ in order to correct for the correlation between spaces, so as to down-weight the individual contribution of strongly correlated CC spaces.

In the ‘explore’ page, measured similarities are shown as filled circles, and inferred similarities as empty ones. Significant similarities (P < 0.05) are shown by large colored circles (filled or empty, correspondingly). Small gray circles are shown otherwise for non-significant similarities. The list of ‘similar’ molecules can be ranked on the basis of the 5 levels of complexity (A-E) by clicking on the level name. By default, we up-rank molecules that are similar to the query molecule in many CC spaces, favoring measured similarities over inferred ones. In the CCweb, we give only the top 125 similar molecules (ensuring that at least 25 similar molecules are selected from each of the 5 CC complexity levels). The user can fetch the *full* list of similar molecules by clicking on the ‘download’ button on the left of the page.

##### Libraries

By default, similar molecules are searched in across the CC (‘all bioactive molecules’). The user can choose to search only in certain chemical collections (approved drugs, LINCS, etc.). Please note that, in contrast to the main page, we explore the complete collection, not only 100 landmark molecules.

#### Statistics and help pages

The user can find summary statistics of the CC, plotted as a slideshow in the statistics page. There is also a help page with a short explanation of the resource and a few FAQ. Links to these pages are placed in the black footer of the home page.

#### Downloads and RESTful acces

CC signatures can be downloaded in HDF5 format (‘download’ link in the home page footer). We also provide programmatic access to our signatures through a REST API. By default, we provide type II signatures. Type I signatures are available upon request.

### Compound collections

We downloaded the following chemical collections from ZINC^87^ (http://zinc15.docking.org, January 2018): approved drugs (‘dbap’), experimental (‘dbex’) and investigational (‘dbin’) drugs, human metabolites (‘hmdbendo’), traditional Chinese medicines (‘tcmnp’), LINCS compounds (‘lincs’), Prestwick Chemical Library (‘pwck’), NIH clinical collection (‘nihcc’), NCI diversity collection (‘ncidiv’), and tool compounds (‘tools’). SMILES strings were converted to InChIKeys using the standardization procedure. When an InChIKey was not explicitly present in the CC, we attempted to match the connectivity layer (i.e. first 14 characters of the key). Unmatched molecules were discarded. Experimental and investigational drugs were merged into the ‘experimental drugs’ (EXD) collection, excluding compounds present in the APD set.

### Transcriptional reversion of familial AD mutations in SH-SY5Y cells

#### Computational screening

##### Proof of principle: correlation between cancer cell line sensitivity and gene expression reversion

We downloaded cell sensitivity data from the GDSC^24^. We mapped 96 of the GDSC drugs on the gene expression dataset (D1) of the CC. Basal gene expression levels of CCLs were converted to z-scores based on gene expression values across the panel^88^. We took the top-250 over- and under-expressed genes for each cell line, according to the expression z-score. To measure ‘reversion’ the direction (up/down) of these two gene sets was flipped.

Cancer cell line transcriptional signatures were then converted to the CC D1 format using the same procedure as that applied to drug signatures, i.e. a two-way GSEA of the signatures was done against Touchstone signatures, results were aggregated over Touchstone cell lines, and a type I signature was eventually obtained. The signature reversion potential of drugs was calculated by simply measuring the distance between type I signatures. Signatures having a Pearson’s correlation > 0.1 between them and their reversed (flipped) version were excluded from the analysis, i.e. they were considered to map poorly to the transcriptional landscape of the CC.

##### Mutated-vs-WT gene expression signatures

AD-specific differential gene expression signatures were obtained by comparing the basal gene expression profiles of APP/PSEN1 mutated with WT SH-SY5Y cells (see below). We generated up/down-regulated gene sets conservatively (adjusted P-value < 0.01, log2-FC > +/- 1.5) and more permissively (P < 0.01, t > +/- 2). Additional versions of the signatures were obtained by keeping only genes related to AD and Tau pathology in OpenTargets (confidence scores of 0.5 (high), 0.2 (medium) and 0.1 (low)). Finally, composite signatures were also derived by simply measuring intersection or union of gene sets (e.g. consensus PSEN1^M146V^ signatures could be obtained from the homozygous and heterozygous clones; see below) (Data S1).

##### Signature reversion

All the signatures above were flipped and converted to the D1 format. Reversion potential (connectivity) of CC compounds was then measured by distance of type I signatures to each AD-related signature. Connectivity scores were robustly normalized (median and median absolute deviation), and aggregated when necessary with the tertile statistic as indicated in^21^. Full results are given in Data S1.

#### CRISPR/Cas9 gene edition for AD cell models

sgRNAs sequences targeting APP and PSEN1 were designed using the Zhang laboratory CRISPR design tool (http://crispr.mit.edu), and cloned into a modified version of pX330 plasmid expressing GFP and puromycin resistance^89^. Next, 200 long single stranded donor oligonucleotides (ssODN) were used as a template for inducing homology-directed repair (HDR) and designed to introduce the desired mutation together with silent mutations to protect both the ssODN template and also the mutated allele once homologous recombination has taken place (Figure S8A). ssODN were purchased from ITD with phosphodiester modification in the 3’. All sequences are listed in Table S2.

SH-SY5Y cells were cultured in DMEM/F12 (1:1) medium supplemented with 10% FBS, glutamine and antibiotics (Thermo Fisher Scientific). For transfection, the SH-SY5Y cells were seeded in T-75 flasks and allowed to grow to 80% confluency. A mixture of X330 plasmid and ssODN template was transfected using linear polyethylenimine (PEI; Polysciences) in Opti-MEM medium (Thermo Fisher Scientific) supplemented with 10% FBS. Three days after transfection cells were trypsinized and seeded again in the presence of 2 µg/ml puromycin (Sigma-Aldrich). Selection pressure with puromycin was kept for 1 week, and then selected cells were allowed to expand and recover for 1-2 weeks. Some of the cells were then used to measure overall HDR efficiency and the rest were single-cell cloned in 96-well plates using a FACSAria II flow cytometer (BD Biosciences). Three to four weeks after cloning, confluent wells were split into two 96-well plates, one to expand the clone and the other to analyze the genotype. DNA extraction was performed adding 50 µl of DirectPCR-tail lysis reagent (VWR) supplemented with 0.4 mg/ml of proteinase K (Roche), and plates were incubated overnight at 55°C. The next day, lysates were moved to 96-well PCR plates and we inactivated Proteinase-K incubating at 85°C for 40 min. Next, 5 µl of lysate was used to amplify by PCR the genomic region surrounding the edition target with recombinant Taq DNA polymerase (Thermo Fisher Scientific), followed by digestion with restriction enzymes in order to screen for the introduction of novel restriction sites encoded in the ssODN template (Figure S8). Using either genomic DNA (gDNA primers) or reverse transcribed RNA (cDNA primers) as template, mutated cells were routinely tested for the presence of the mutation by selective digestion with restriction enzymes (Figure S8B) of PCR-amplified DNA fragments surrounding the mutation. All primer sequences are listed in Table S2. All restriction enzymes were purchased from New England Biolabs.

Of the clones isolated, we obtained homozygous clones (APP^V717F/V717F^ and PSEN1^M146V/M146V^) and also heterozygous mutants in which the second allele had a two-nucleotide deletion leading to a displacement in the reading frame and a premature stop codon, therefore called “null” (APP^V717F/null^) or, in the case of the M146V mutation, a three-nucleotide deletion encoding for an amino acid deletion at position 149 (PSEN1^M146V/L149Δ^). We then measured the two main forms of Aβ peptide secretion (Aβ42 and Aβ40) in all the isolated clones, and we observed an increase in the Aβ42/Aβ40 ratio (Figure S8C).

#### Aβ quantification

Aβ peptides were quantified by ELISA-based assays using either the 6E10 Aβ Triplex by MesoScale Diagnostics or the Wako ELISA kit Human β Amyloid (1-40) and Wako ELISA kit Human β Amyloid (1-42) High-Sensitivity, following manufacturer’s instructions. Direct comparison of the results showed similar results for the two quantification assays.

#### Drug treatment of SH-SY5Y clones

Cells were differentiated for 3 days in neurobasal medium supplemented with B27, glutamax (all Thermo Fisher Scientific), 10 µM retinoic acid (Sigma-Aldrich) and 50 ng/mL Brain-Derived Neurotrophic Factor (BDNF; Peprotech). Then, medium was renewed in the presence of the indicated concentration of drugs. All drugs were dissolved in DMSO, and controls of cells treated with DMSO were run in parallel, at a final concentration of 0.1% DMSO. After 3 days, supernatants were stored at −80°C for Aβ measurement and cells were either incubated for 1 h in the presence of 3-(4,5-dimethylthiazol-2-yl)-2,5-diphenyltetrazolium bromide (MTT) and lysed in DMSO to check the viability, or lysed with RTL buffer (Qiagen) for RNA extraction using the RNAeasy mini kit (Qiagen). Three independent experiments were performed.

#### Gene expression

To obtain the signature profile derived from these fAD mutated cells, SH-SY5Y WT and mutated cells were differentiated for 6-7 days in the presence of retinoic acid and BDNF to recapitulate a phenotype more similar to neurons as previously described^25^. The secretion of Aβ followed the same pattern as that observed in non-differentiated cells. Samples of purified RNA of WT (APP/PSEN1^WT/WT^), APP^V717F/null^, PSEN1^M146V/L149Δ^ and PSEN1^M146V/M146V^ (clone #2) were extracted and submitted to the IRB Functional Genomics Facility, where sample quality was assessed using an Agilent Bioanalyzer. The whole-genome expression profile was generated using Affymetrix PrimeView arrays. Three independent experiments were used to obtain the expression profiles.

#### Evaluation of results

The ‘reversion capacity’ of tested drugs was measured as follows. We ranked the differential gene expression results of the treated-vs-untreated comparison performed on mutated cells (ranked list 1). In parallel, we ranked the differential gene expression results of the mutated-vs-WT comparison (ranked list 2). In Figure 4B, we simply traverse ranked list 2 (x-axis, from both tails) and measure the identification of genes at the other end (top-250) of ranked list 1 (y-axis). Ranked list 1 was randomized to assign significance to observations. In order to obtain a ‘reversion strength’ value per gene, we designed a reversion score based on the difference in ranks between mutated-vs-WT (reference) and treated-vs-control gene expression profiles. We scaled reversion scores, ranging from −1 (under-expressed genes in the reference are over-expressed upon treatment) to +1 (viceversa). To test whether reversed genes were enriched in AD genes (OpenTargets score > 0.5), we performed a weighted Kolmogorov-Smirnov, taking as weights the absolute value of the reversion score^90^.

### Identification of small molecule mimetics of biodrugs against IL2R, IL-12 and EGFR

#### Computational screening

##### Biodrug-related signatures

Biodrugs were defined by their targets (i.e. IL2R, IL12B and EGFR). We derived D1 (transcriptional), C3 (pathway), C4 (biological process) and C5 (interactome) CC signatures for these three targets. C3 and C4 signatures were obtained by simply mapping pathways and biological processes of the targets, respectively, and expressing them as CC signatures by TF-IDF/LSI transformation. Regarding C5 signatures, we applied the same procedure than the one applied to compounds, i.e. we mapped target neighbors in interactomes using HotNet2 and then we obtained the corresponding type I signature. For D1, we downloaded gene expression signatures from shRNA experiments obtained from LINCS L1000, and mapped them analogously to small molecule perturbations.

##### Matching biodrug-related signatures

We devised a computational screening for drugs in D1 spaces and in any of the C3-5 spaces. We asked for candidates to be amongst the top 250 drugs in terms of similarity of the target signature to at least one of the C3-5 spaces, and ranked them on the basis of similarity of transcriptional profiles (mimicking, i.e. D1 similarity).

#### Cells

PBMC were purchased from StemCell Technologies and maintained in RPMI medium supplemented with 10% FBS, glutamine and antibiotics (Thermo Fisher Scientific). Pre-stimulated PBMC were obtained by culturing 10^6^ PBMC/mL with 0.5 µg/ml soluble anti-CD28 (CD28.2) and anti-CD3 (OKT3) antibodies (Thermo Fisher Scientific) for three days. After this stimulation period, cells were washed and left untreated for three days before re-stimulation. H1650, Jurkat and MT-4 cells were cultured in RPMI supplemented with 10 % fetal bovine serum (FBS), glutamine and antibiotics (Thermo Fisher Scientific). A431 and HeLa cells were cultured in DMEM supplemented with 10% FBS, glutamine and antibiotics (Thermo Fisher Scientific). NK-92 cells were purchased from ATCC (CRL-2407), cultured in alpha-MEM without ribo- and deoxyribo-nucleosides (Thermo Fisher Scientific), and supplemented with FBS and horse serum (both Thermo Fisher Scientific), 0.2 mM inositol (Sigma-Aldrich), 0.1 mM 2-mercaptoethanol (Sigma-Aldrich), and 0.02 mM folic acid (Sigma-Aldrich), penicillin/streptomycin and glutamine (both Thermo Fisher Scientific). 100 U/ml of recombinant IL-2 (Peprotech) was added every 2-3 days.

#### PBMC proliferation assay

Resting and pre-stimulated PBMC where loaded with 2 µM CFSE (Thermo Fisher Scientific) in PBS with 0.1% FBS for 7 min at 37°C. After two washes in complete medium, PBMC were pre-treated for 1 h with the corresponding drugs, followed by stimulation with 0.5 ng/mL IL-2 or 5 µg/mL PHA (Sigma-Aldrich). Three days after stimulation, cell fluorescence was measured using a Gallios Flow Cytometer (Beckton Coulter). Analysis was performed with FlowJo software.

#### Phospho-STAT5 quantification by flow cytometry

Pre-stimulated PBMC pre-treated for 1 h with the corresponding compounds were stimulated for 20 min with 0.5 ng/mL IL-2, fixed with Fix Buffer I (BD Biosciences), permeabilized with Perm Buffer III (BD Biosciences), and finally stained with a PE-labelled anti-phospho-Stat5 (pY694; BD Biosciences). Staining was measured in a Gallios Flow Cytometer and analysis was performed with FlowJo software. Two compounds, SU11652 and Z55175877 showed autofluorescence at high concentrations when cells were analyzed by flow cytometry, therefore STAT5 phosphorylation was measured by western blot as detailed below.

#### Proliferation of cell lines

5·10^4^ Jurkat or MT-4 cells were incubated in the presence of the indicated compounds. In the case of HeLa cells, 5·10^3^ cells were seeded the day before the compounds were added to the supernatant. After 3 days, cells were incubated for 1 h in the presence of 3-(4,5-dimethylthiazol-2-yl)-2,5-diphenyltetrazolium bromide (MTT) and viability/proliferation was quantified as indicated above.

#### IL-12 stimulation

The day before the experiment, NK-92 cells were counted, washed with RPMI supplemented with 10% FBS and seeded in 24-well plates (3·10^5^ cells/well) in RPMI 10% FBS in the absence of IL-2. On the day of the experiment, cells were pre-incubated with the indicated compounds for 1 h and then stimulated with 50 ng/mL IL-12 (Peprotech). Cells were pelleted after 1 h of stimulation and lysed for western blot analysis or kept up to 5 h in culture. They were then pelleted and RNA was extracted for quantitative PCR analysis as indicated below.

#### Quantitative PCR

For quantitative PCR (qPCR), purified RNA samples were reverse transcribed with the High-Capacity cDNA Reverse Transcription Kit (Thermo Fischer Scientific) and qPCR was performed in a QuantStudio 6 Flex Real-Time PCR System (Thermo Fischer Scientific) using the LightCycler 480 SYBR Green I Master mix (Roche). Ct values were normalized using GAPDH as reference gene and the ΔΔCt method to quantify the fold change of the gene of interest. Primers are shown in Table S2.

#### EGFR analysis

0.15·10^6^ A431 or H1650 cells were seeded in 24-well plates the day before the experiment. Cells were treated with the indicated inhibitors for 24 h. Cells were then washed with PBS and lysed for western blot analysis.

#### Western Blot

Cells stimulated with or without cytokines were washed in PBS, concentrated and resuspended in lysis buffer (50 mM Tris-HCl [pH 7.5], 1 mM EGTA, 1 mM EDTA, 1% [wt/wt] Triton X-100) supplemented with protease inhibitor cocktail (Roche) and phosphatase inhibitor cocktail (Roche). Lysates were subjected to SDS-PAGE in Mini-PROTEAN TGX Stain-Free Precast Gels (Biorad) and transferred to a polyvinylidene difluoride membrane using the Trans-Blot Turbo Transfer System (Biorad). Images of developed blots were acquired with the Chemidoc Touch Imaging System (Biorad).

The following antibodies were used for immunoblotting: horseradish peroxidase-conjugated secondary antibodies (Thermo Fisher Scientific), anti-Actin (Merck), anti-Stat5 (D206Y; Cell Signaling Technology) and anti-Stat4 (C46B10; Cell Signaling Technology). Phosphospecific antibodies recognizing phospho-Tyr693 of Stat4 and phospho-Tyr694 of Stat5 (D47E7), were also from Cell Signaling Technology. The EGFR monoclonal antibody (528) was purchased from Thermo Fisher Scientific.

#### Statistical Analysis

Data were analyzed with the Prism statistical package. Unless otherwise indicated in the figure legend, *P-*values were calculated using an unpaired, one-tailed, Student *t*-test.

## Acknowledgements

We would like to thank the SB&NB lab members for their support and helpful discussions. We are grateful to the Broad Institute and National Center for Advancing Translational Sciences (NCATS-NIH) for providing compounds upon request. We also thank the IRB Barcelona Biostatistics and Bioinformatics Unit and the IRB Functional Genomics Facility, and J. Duran-Frigola for the website design. P.A. acknowledges the support of the Spanish Ministerio de Economía y Competitividad (BIO2016-77038-R) and the European Research Council (SysPharmAD: 614944).

## Author contributions

M.D-F., E.P. and P.A. designed the study, analyzed the results and wrote the manuscript. M.D-F. did the computational analysis, together with M.B., T.J-B. and D.A. O.G-P. implemented the web-server. E.P. and V.A. carried out the experimental validations. All authors have read and approved the manuscript.

## Supplemental Information

**Data S1.** *Reversion of transcriptional signatures of familial Alzheimer’s disease (fAD) mutations.* This dataset contains the results of the search for LINCS molecules that might *revert* transcriptional signatures of fAD mutations in SH-SY5Y cells, and the analyses of transcriptional signatures for the tested compounds. We first provide the signatures obtained by differential gene expression measurements of PSEN1 or APP mutated cells relative to WT, and also the additional signatures obtained as indicated in Methods (see *Computational legend* sheet for details). Second, we provide the connectivity (reversion) score of all the previous signatures for each compound in D1 space. Next, we provide the results of the experimental gene expression reversion with the tested compounds, namely Noscapine, Palbociclib, and AG-494 (see the *Reversion legend* sheet for details). In particular, we analyze the best-reversed genes with a normalized score ranging from 1 (an up-regulated gene that is highly down-regulated upon treatment) to −1 (a down-regulated gene that is highly up-regulated upon treatment). Finally, we provide the vendors for the commercial compounds used in this section.

**Data S2.** *Small molecule analogs of biodrugs.* This dataset contains the signature ‘matching’ search to identify small molecule drugs that might resemble biodrugs against IL-2 receptor, IL-12 and EGF receptor. Overall, we provide a score for the mimicking of transcriptional signatures obtained from silencing experiments (D1), matching of pathways (C3), biological processes (C4) and network environments (C5) related to the biodrug target, and a P-value and a rank for all the comparisons (see the *Legend* sheet for details). Finally, we provide the vendors for the commercial compounds used in this section.

---

**Figure S1.**
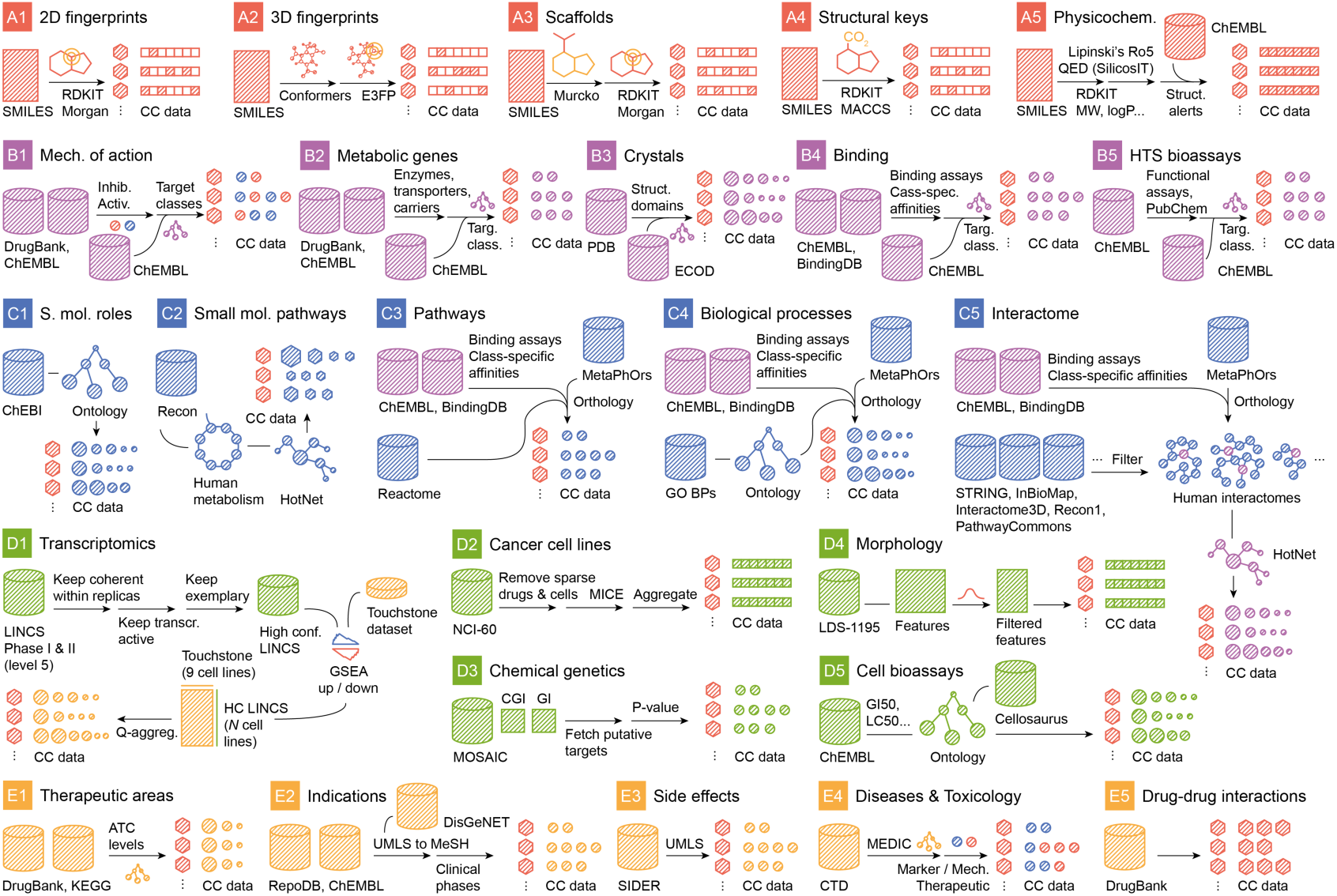
Data pre-processing. Schematic representation of pre-processing pipelines for each of the 25 CC spaces. Details are given in the Methods.

**Figure S2.**
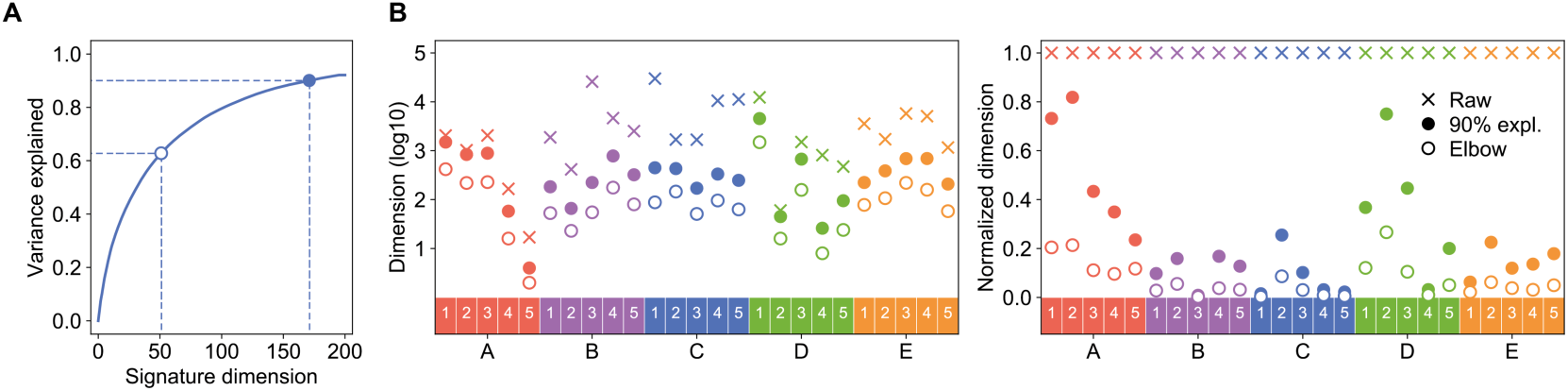
Dimensionality reduction (type I signature). (A) An illustrative scree plot (C3: Signaling pathways), showing the dimension that keeps 90% of the variance (bold dot) and the elbow of the curve (white dot). (B) In the left panel, number of dimensions of the signature type I (bold dot), the elbow point (white dot) and original (raw) dimensions (cross). In the right panel, values are normalized by the original dimensions. Original dimensions correspond to the size of the feature space of the raw data (e.g. number of targets, number of bits in a 2D fingerprint, number of measured genes, number of morphological features, number of yeast mutants, etc.).

**Figure S3.**
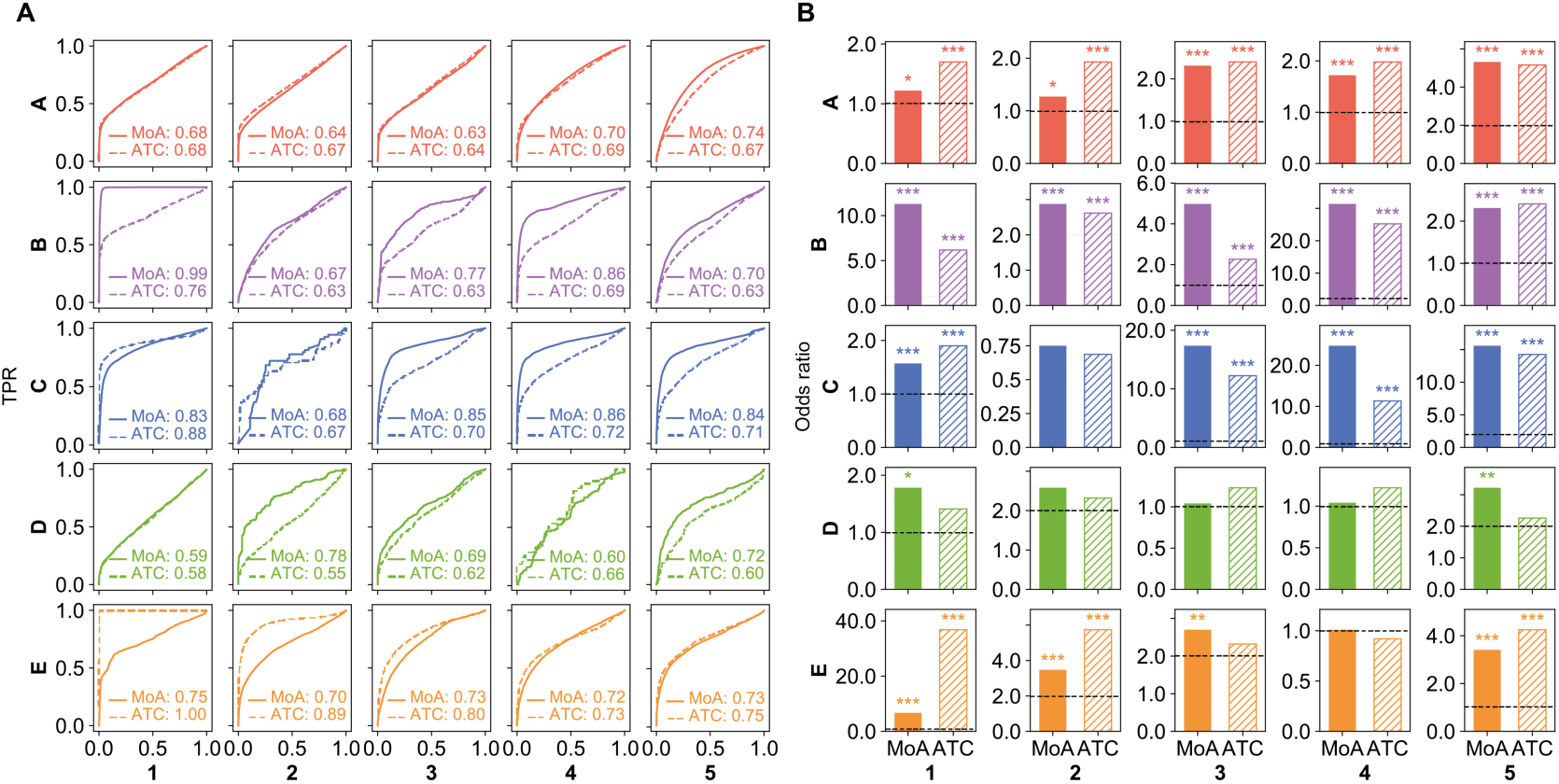
MoA and ATC validations. (A) ROC curves measuring the association between type I signature cosine similarities and the fact that two drugs share a MoA or an ATC code (level 3). (B) Likewise, odds ratio and significance (*** P < 0.0001, ** P < 0.001 and * P < 0.01) of a Fisher’s exact test performed on a contingency table classifying drug pairs as having the same MoA/ATC code or not, and as belonging to the same cluster or not (see Methods).

**Figure S4.**
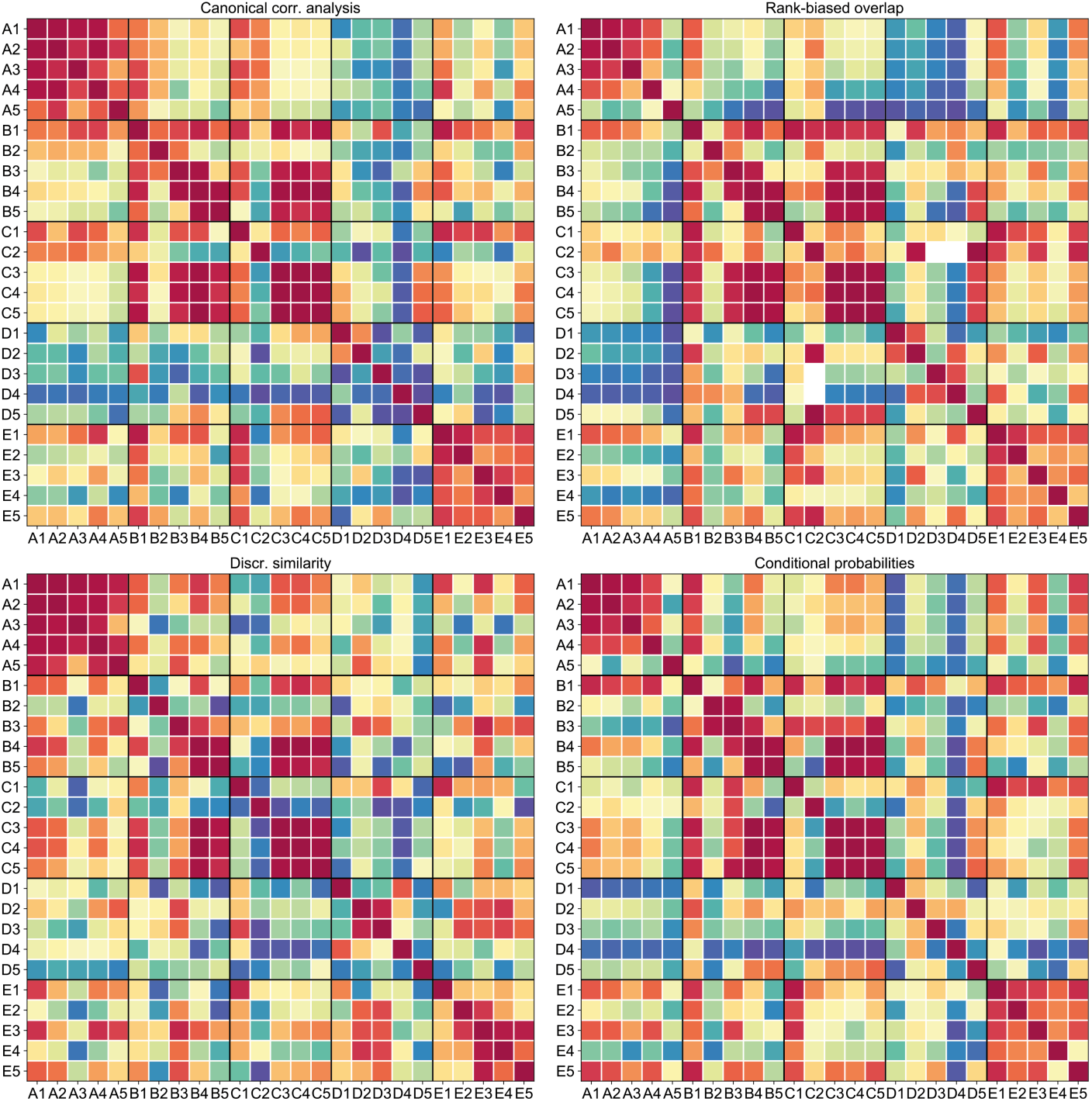
Correlations between CC spaces. Four dataset correlation measures are shown, as described in the Methods.

**Figure S5.**
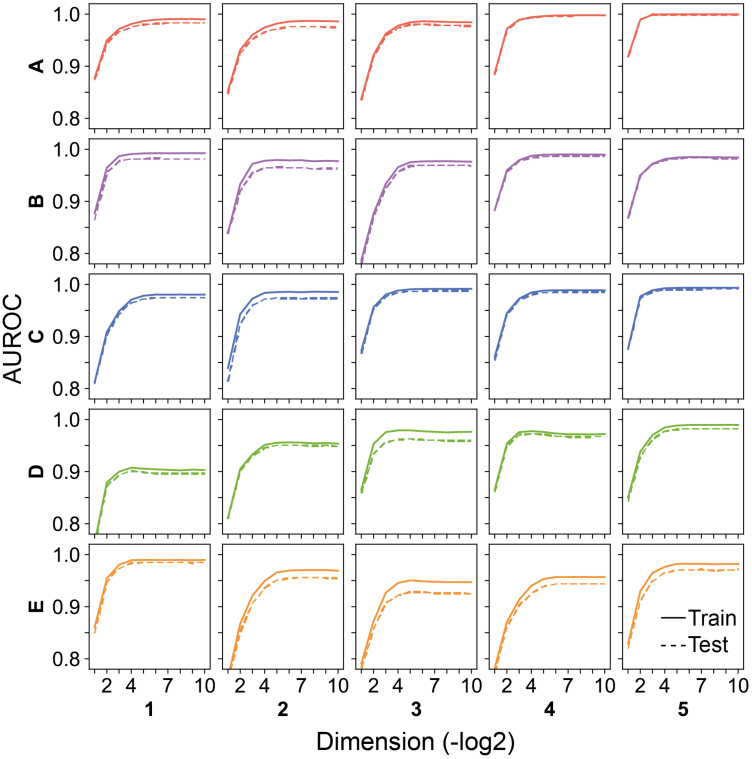
Similarity network embedding (type II signature). ROC curves of ‘link prediction’ exercises, performed by removing 20% of the similarity links (P < 0.01). Performance is evaluated at different embedding dimensions.

**Figure S6.**
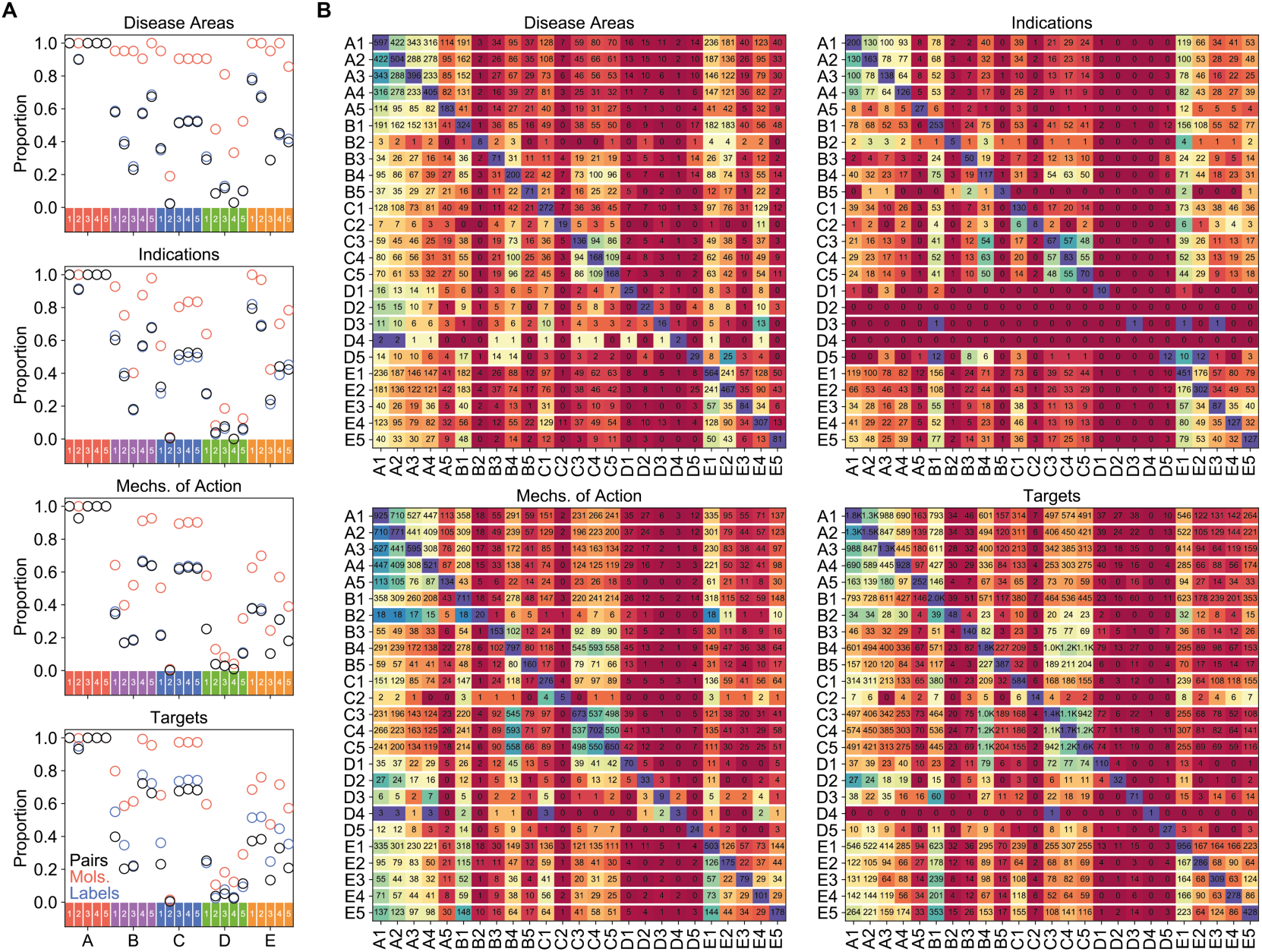
Similarity-based label prediction. (A) Proportion of molecule-label pairs (black), molecules (red) and labels (blue) that can be considered for prediction in each of the CC spaces. We require a label to be annotated with at least 5 molecules in a given CC space. (B) Overlap between true positives across CC spaces. The matrix is read row-wise, e.g. 191 of the 597 true positives obtained by A1 are also found by B1. The color scale is normalized to the diagonal in each row.

**Figure S7.**
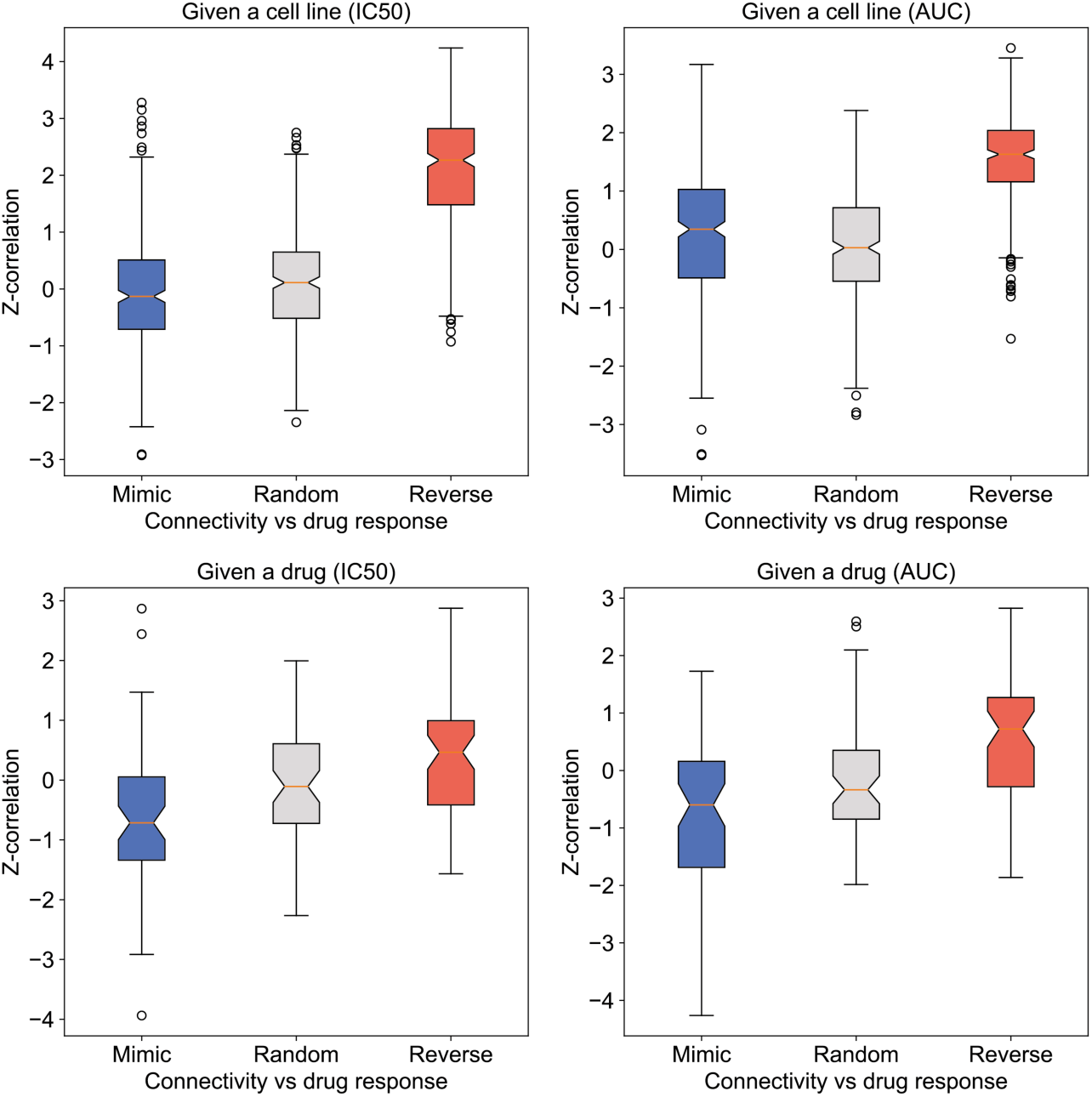
Correlation between GDSC drug sensitivity and signature reversion scores. This plot certifies that transcriptional signature reversion with CC signatures performs as expected with cancer cell lines, i.e. drugs that more strongly reverse transcriptional traits of certain cancer cell lines are indeed more potent against these cells. Upper panels rank, for each cell line, drugs according to their ability to mimic (blue) or revert (red) basal gene expression profiles of the cells, based on D1 CC data. These ‘connectivity’ scores are correlated to cell sensitivity (expressed as IC50 or AUC of the response curve). Likewise, lower panels rank, for each drug, cell lines according to their basal gene expression ‘connectivity’ to the drug signature. The Fisher z-transformation is applied to correlation scores to correct for the different number of cell lines per drug and drugs per cell line.

**Figure S8.**
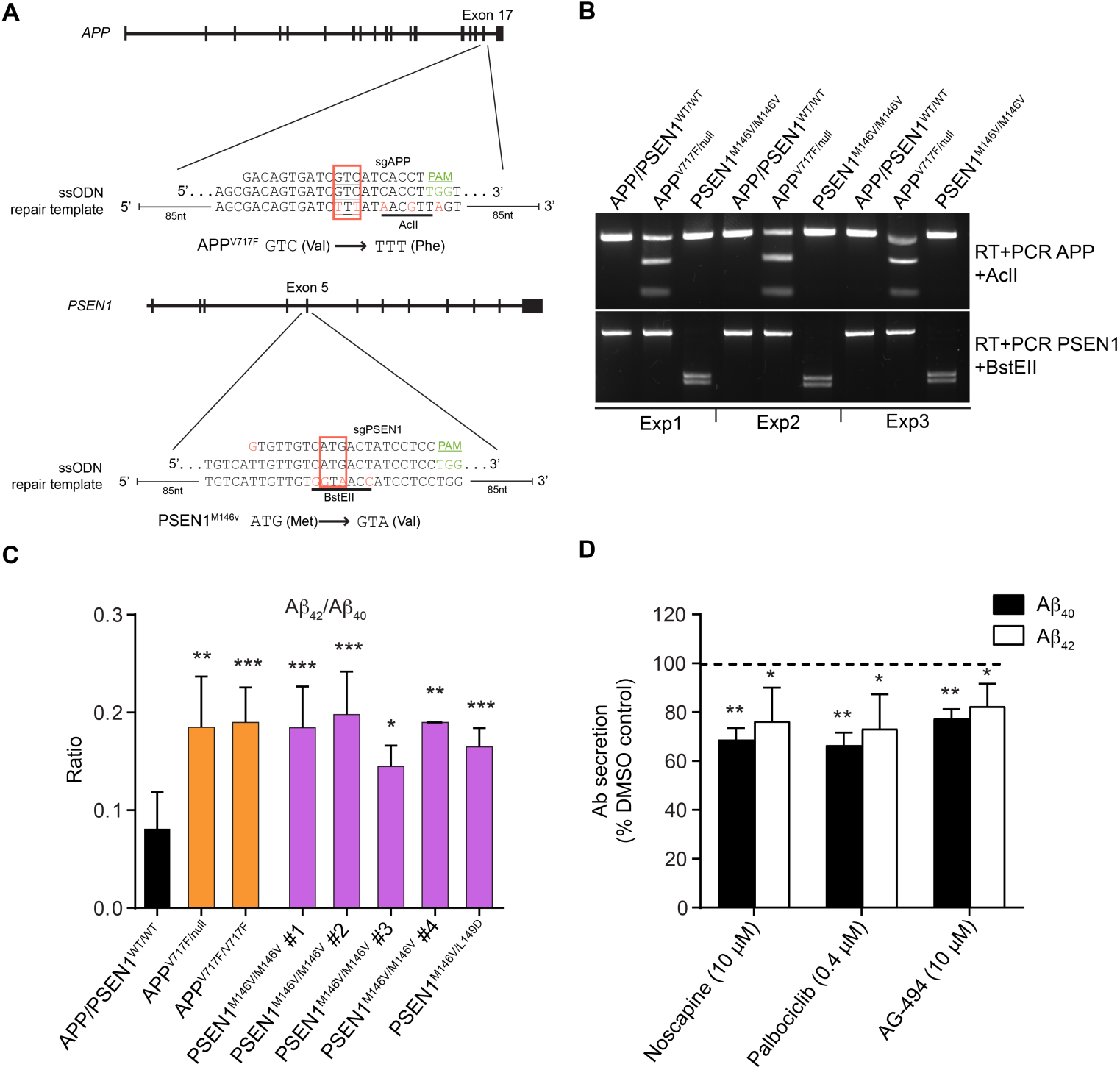
CRISPR/Cas9-mediated generation of SH-SY5Y cells expressing fAD mutations. (A) Strategy designed to introduce APP^V717F^ and PSEN1^M146V^ mutations in the genome of SH-SY5Y cells. Guide RNA sequence (sgRNA) and single-stranded DNA template are indicated. Silent mutations are introduced to protect the mutated alleles from new CRISPR/Cas9-mediated cuts and also to introduce restriction sites to identify the mutated alleles. Sequences can be found in Table S2. (B) Mutated cells can be identified by digestion with the indicated restriction enzymes after reverse transcription followed by PCR amplification of the PSEN1 or APP mutated regions. Clones were routinely tested to confirm their genotype. Representative gels are shown. (C) Independent APP^V717F^ and PSEN1^M146V^ mutant clones have significantly increased Aβ_42_/Aβ_40_ ratios compared with control wild-type SH-SY5Y cells, reflecting a relative increase in the generation of Aβ_42_. Mean ± SD of 2-5 independent experiments are shown. All clones were compared with APP-PSEN1 WT cells (APP-PSEN1^WT/WT^). (D) Normalized Aβ_40_ and Aβ_42_ secretion in differentiated WT SH-SY5Y cells treated with the indicated compounds. Mean ± SD of three independent experiments are shown. One-sample t-test comparing column means to the reference value of 100. *** P < 0.001, ** P < 0.01, * P < 0.05.

**Figure S9.**
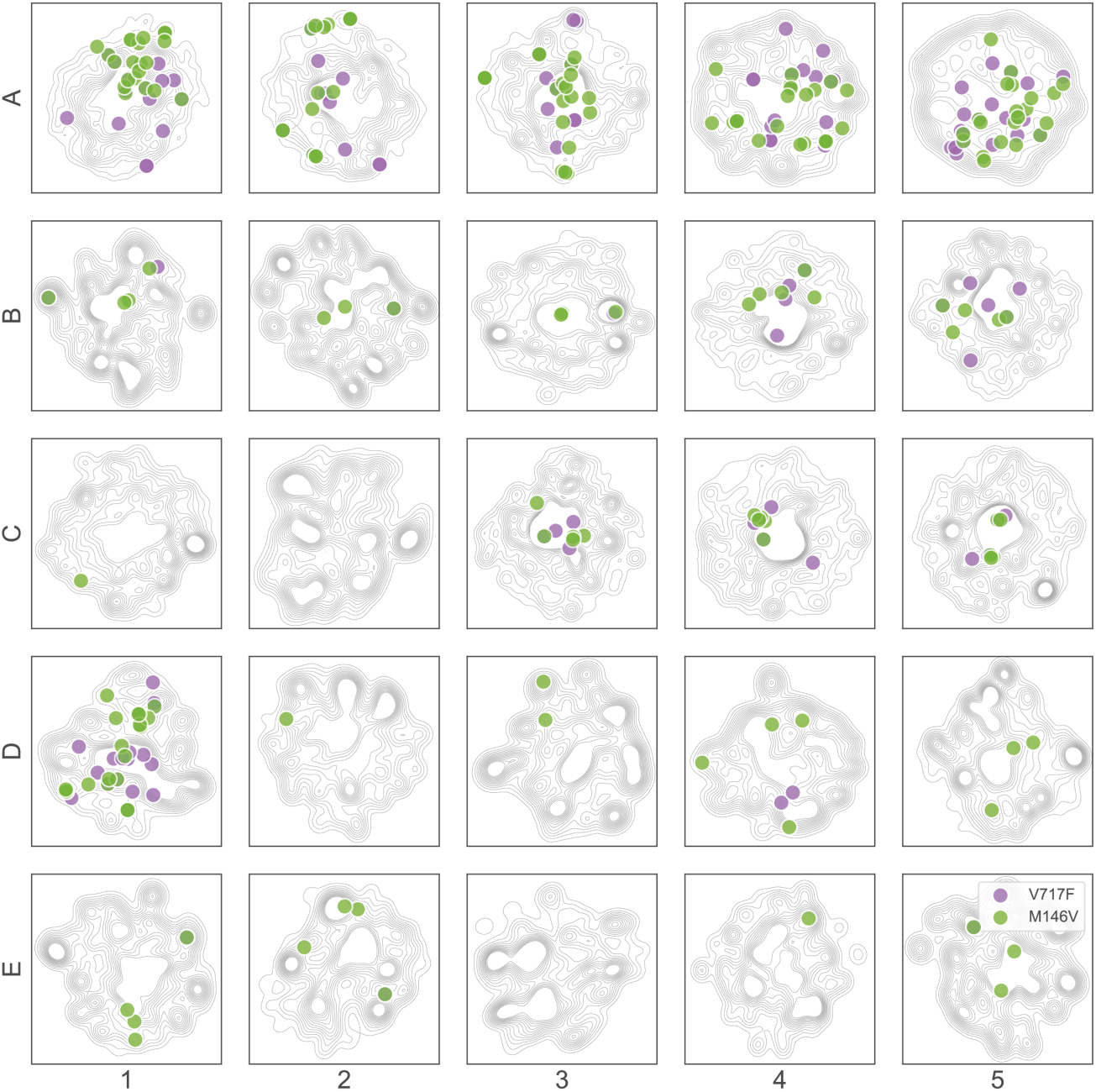
Experimentally tested AD candidate compounds, mapped to the CC. Candidates for signature reversion in the AD signature reversion experiment are located in 2D projections of the CC. In purple, we show the candidates for the APP^V717F^ mutant, and in green the candidates for the PSEN1^M146V^ mutant.

**Figure S10.**
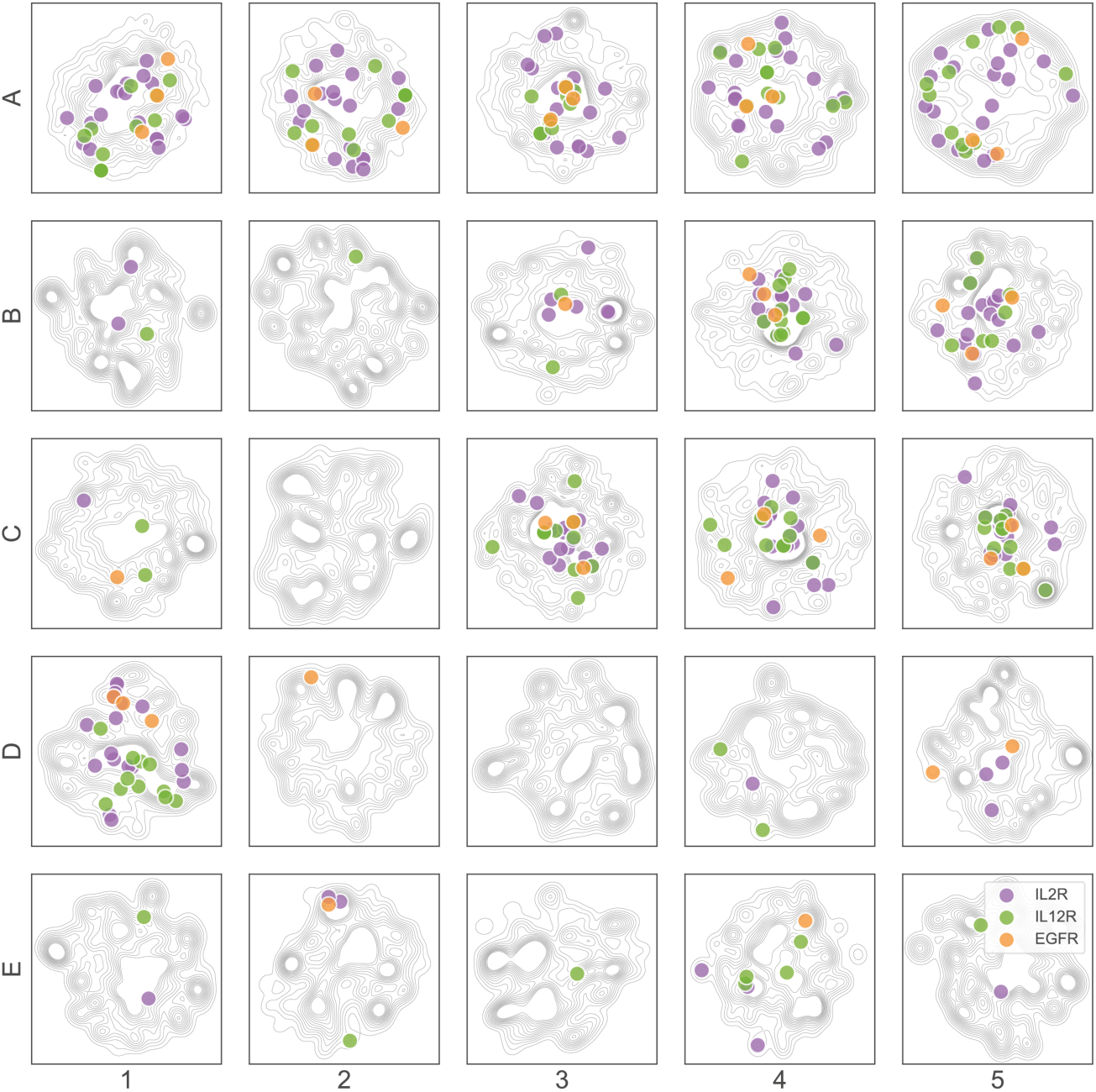
Experimentally tested biodrug mimetics. Candidates selected in the IL2R (purple), IL-12 (green) and EGFR (orange) computational screening are located in 2D projections of the CC.

**Figure S11.**
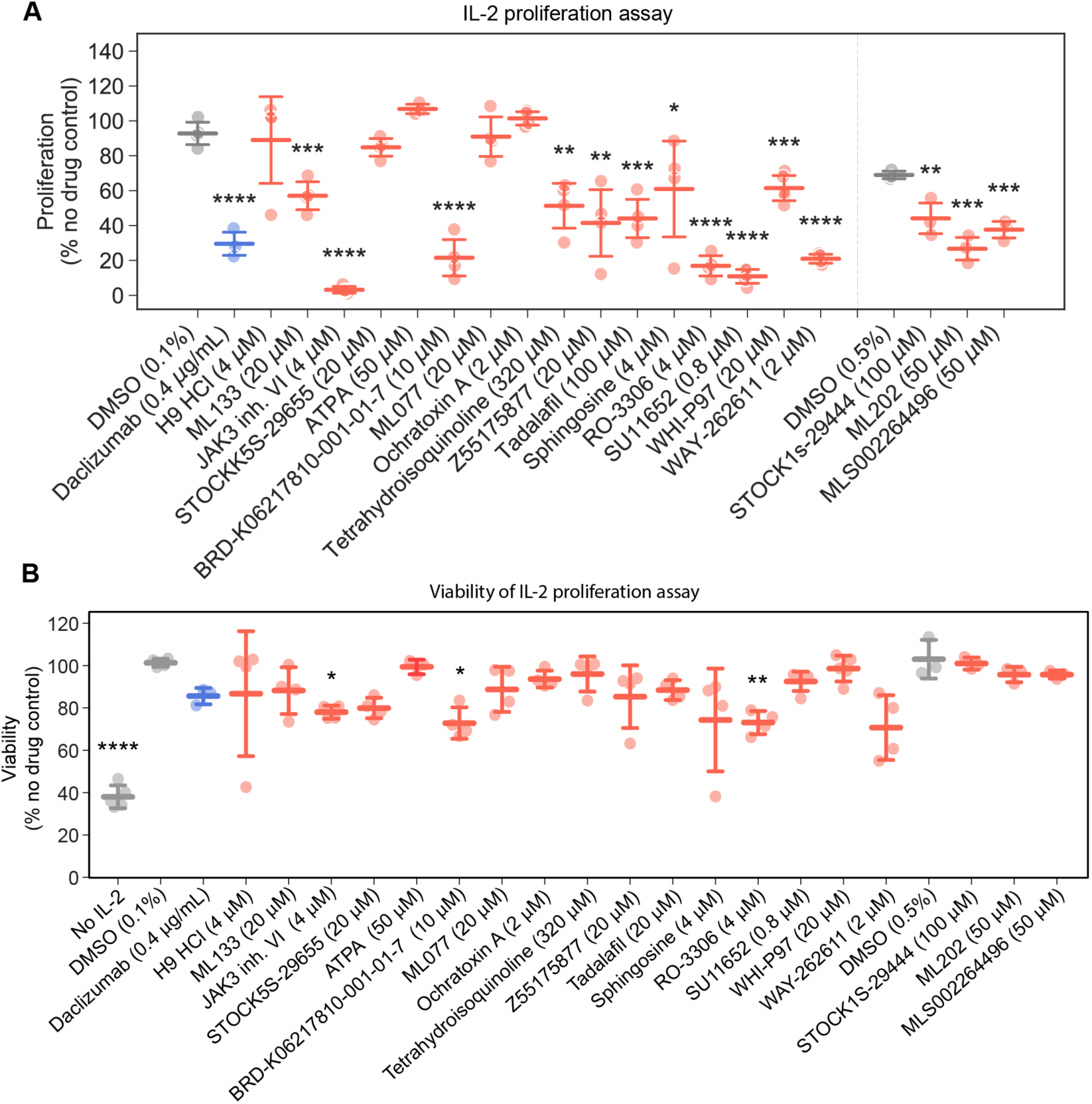
Proliferation and viability of pre-stimulated PBMC treated with IL-2. (A) CD3/CD28 pre-stimulated PBMC were left without treatment for 3 days, labelled with CFSE and then stimulated with IL-2 (0.5 ng/mL). Three days after stimulation, proliferation was measured by flow cytometry as CFSE label decay. Values were compared with the corresponding vehicle controls (DMSO 0.1% for the left panel and DMSO 0.5% for the right panel). Mean ± SD of 3-5 independent experiments are shown. (B) CD3/CD28 pre-stimulated PBMC were left untreated for 3 days and then stimulated with IL-2. Three days after stimulation, proliferation was measured by flow cytometry and viability was quantified measuring the percentage of events detected in the gate corresponding to living cells. Values were normalized as percentage of cells stimulated in the absence of drug. Mean ± SD of 3-5 independent experiments are shown. Because lack of IL-2 signaling may affect cellular viability, values were compared with the sample treated with daclizumab at 0.4 µg/ml. **** P < 0.0001, *** P < 0.001, ** P < 0.01, * P < 0.05.

**Figure S12.**
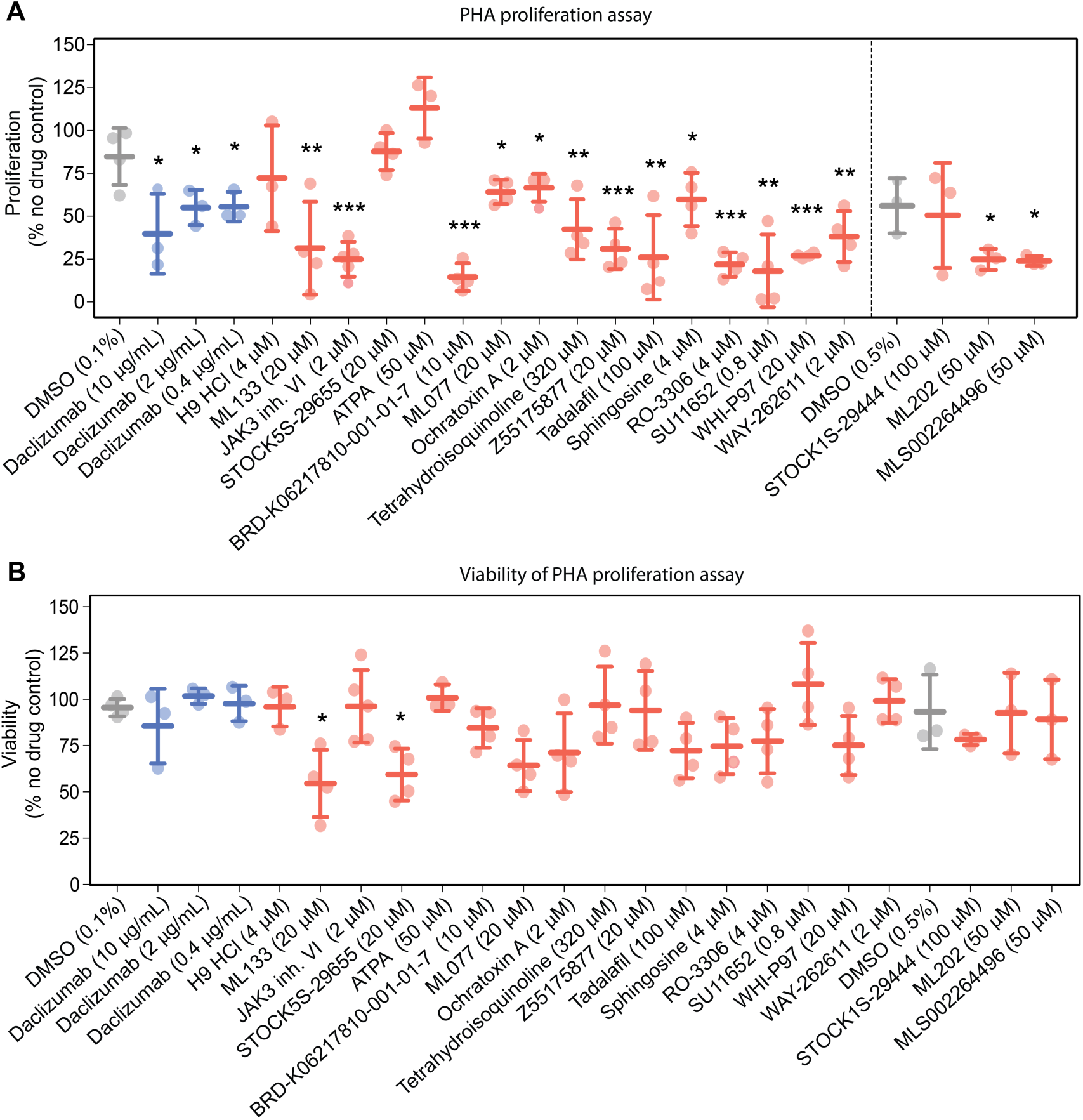
Proliferation and viability of resting PBMC treated with PHA. (A) Resting PBMC were labelled with CFSE and stimulated with PHA. CFSE decay was assessed by flow cytometry three days later. Cell proliferation was quantified as the percentage of cells with reduced CFSE compared with non-stimulated lymphocytes. Values were normalized as percentage of proliferation compared to cells stimulated in the absence of drug. Values were compared with the corresponding vehicle controls (DMSO 0.1% for the left panel and DMSO 0.5% for the right panel). (B) In the same experiment described in (A), viability was quantified by measuring the percentage of events detected in the gate corresponding to living cells. Values were normalized as percentage of viability compared with cells stimulated in the absence of drug. Mean ± SD of 3-5 independent experiments are shown. Because lack of IL-2 signaling may affect cellular viability, values were compared with the sample treated with daclizumab at 10 µg/ml. **** P < 0.0001, *** P < 0.001, ** P < 0.01, * P < 0.05.

**Figure S13.**
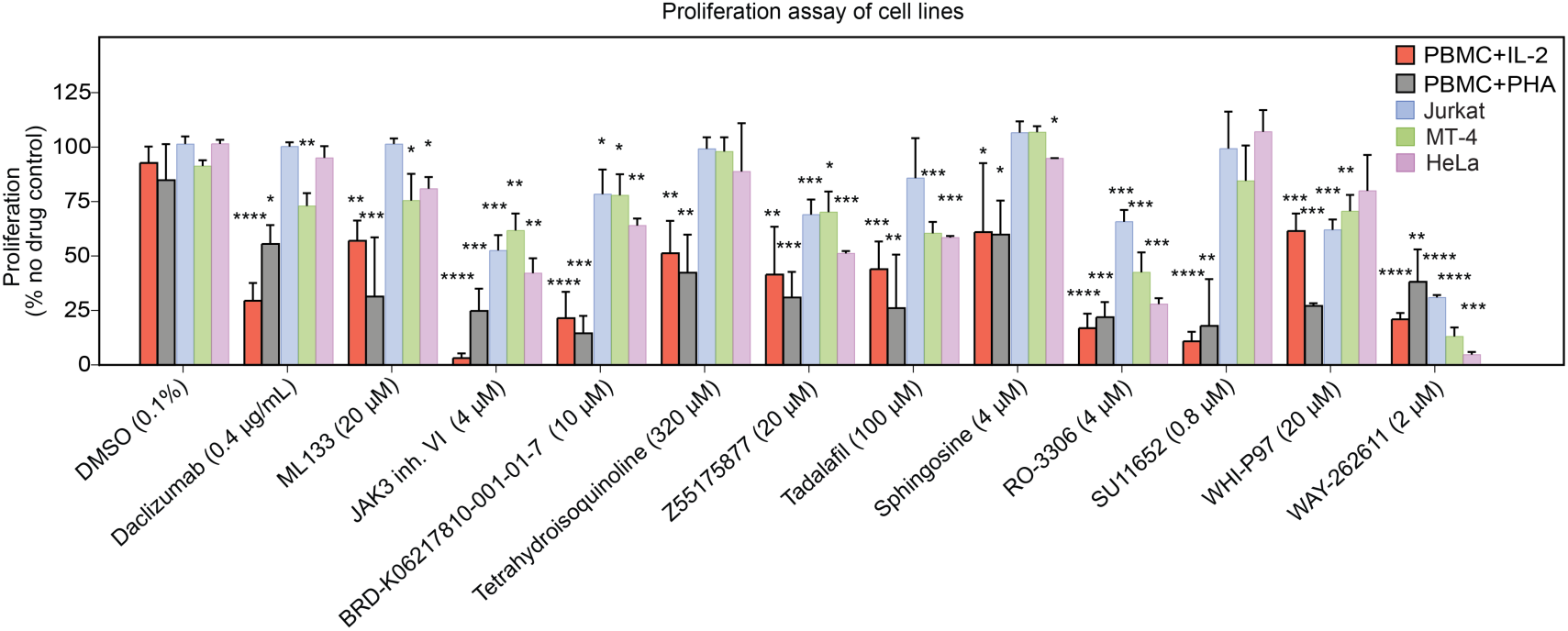
Proliferation of cell lines. Proliferation of PBMC stimulated with IL-2 (red) or PHA (grey), as shown in Figure S11A and S12A, respectively, are depicted together with the proliferation of cancer cell lines Jurkat (orange), MT-4 (blue) and HeLa (green). Cells were treated for 72 h with the indicated drugs, and proliferation was measured by the MTT assay. Values were normalized as percentage of cells compared with cells left in the absence of drug. Values were compared with the vehicle control (DMSO 0.1%). Mean ± SD of 2-3 independent experiments are shown. **** P < 0.0001, *** P < 0.001, ** P < 0.01, * P < 0.05.

**Figure S14.**
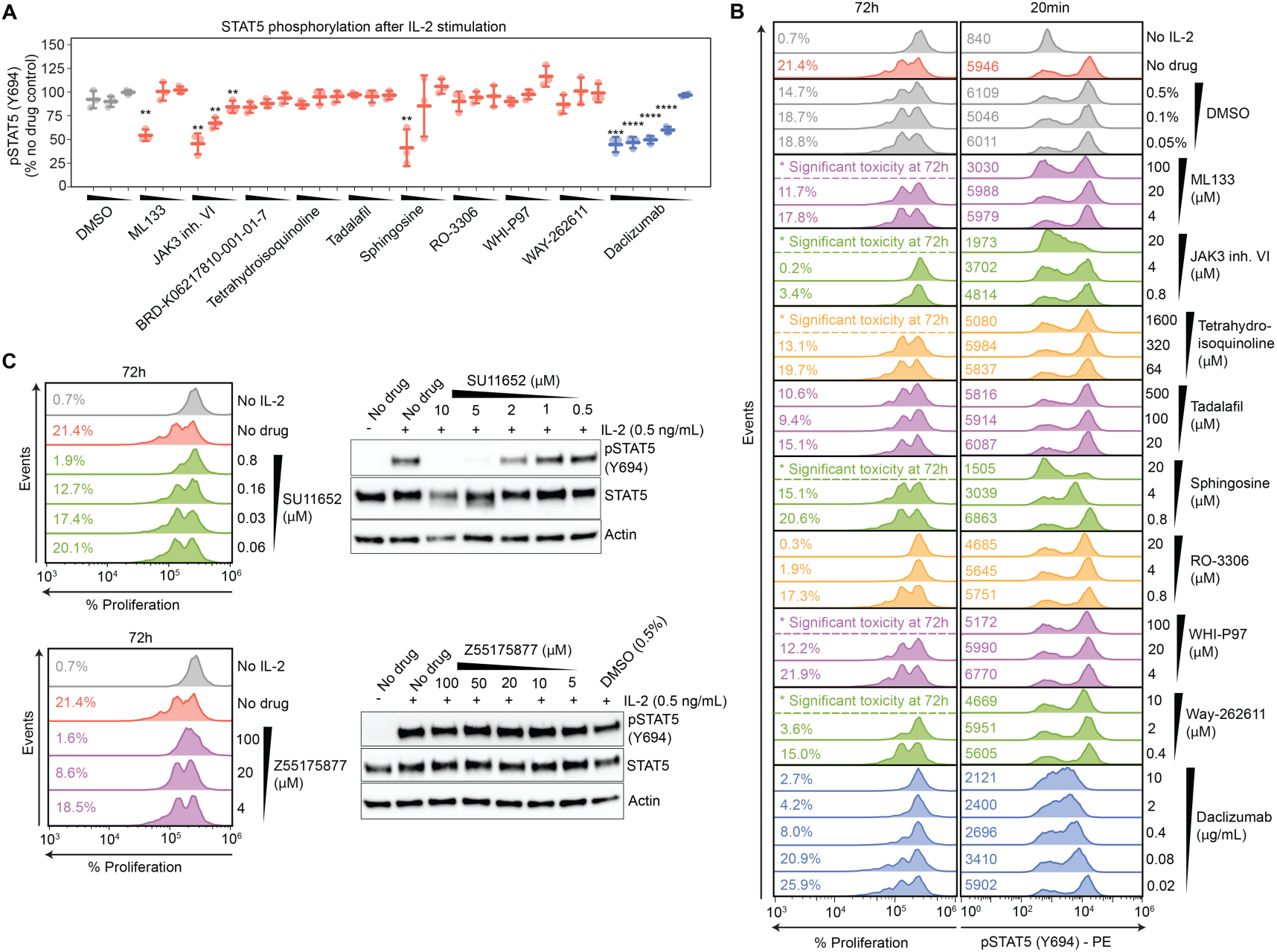
Phosphorylation of STAT5. (A) Pooled results of STAT5 phosphorylation quantification by flow cytometry. Pre-stimulated PBMC were pretreated for 1 h with the indicated doses of compound, and cells were stimulated with IL-2 (0.5 μg/mL) for 20 min. Cells were fixed, and STAT5 phosphorylation was measured by flow cytometry. Values were normalized comparing with cells stimulated in the absence of drug. Mean ± SD of 2-3 independent experiments are shown. **** P < 0.0001, *** P < 0.001, ** P < 0.01, * P < 0.05. (B) Representative flow cytometry results for selected compounds, showing PBMC proliferation (after 72 hours) and STAT5 phosphorylation (after 20 minutes) following IL-2 stimulation. We plot the percentage of cells that have proliferated and the mean fluorescence values corresponding to the phospho-STAT5 staining. A representative experiment, out of 3 replicates, is shown. (C) Representative results for compounds that showed autofluorescence in flow cytometry analysis of STAT5 phosphorylation. Cell proliferation was quantified by flow cytometry and STAT5 phosphorylation was measured by western blot. Representative blots out of 3 independent experiments are shown. Compounds are listed in Table S1.

**Table S1.**
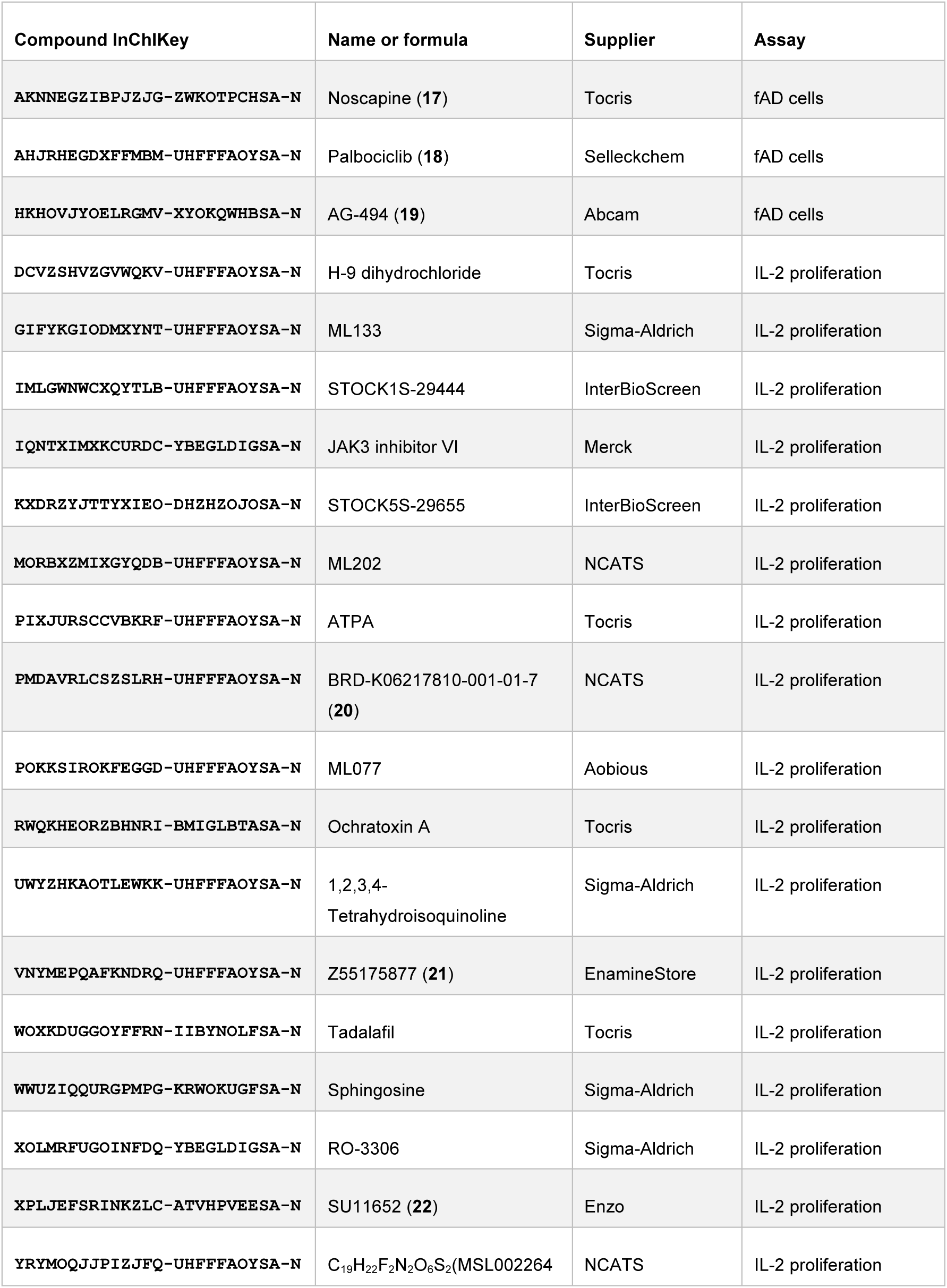
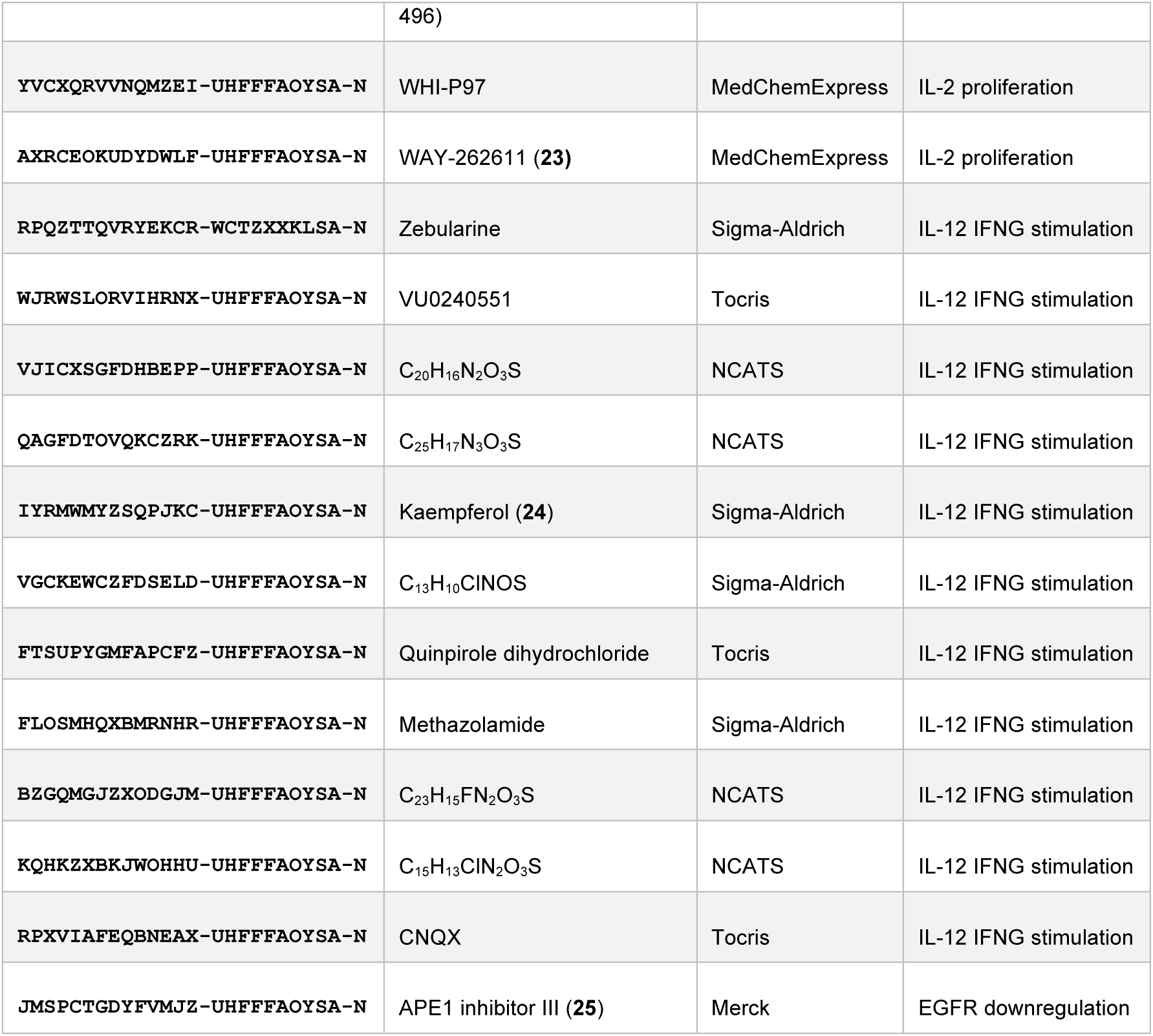
Compounds used in the signature reversion predictions. Drugs were purchased from the indicated suppliers or kindly provided by The Broad Institute or the National Center for Advancing Translational Sciences (NCATS) of the National Institutes of Health (US). CD25/IL-2 R alpha antibody (daclizumab) was purchased from Novus Biologicals. APE1 inhibitor III is an analog of the APE1 inhibitor initially identified in the computational screening.

**Table S2.**
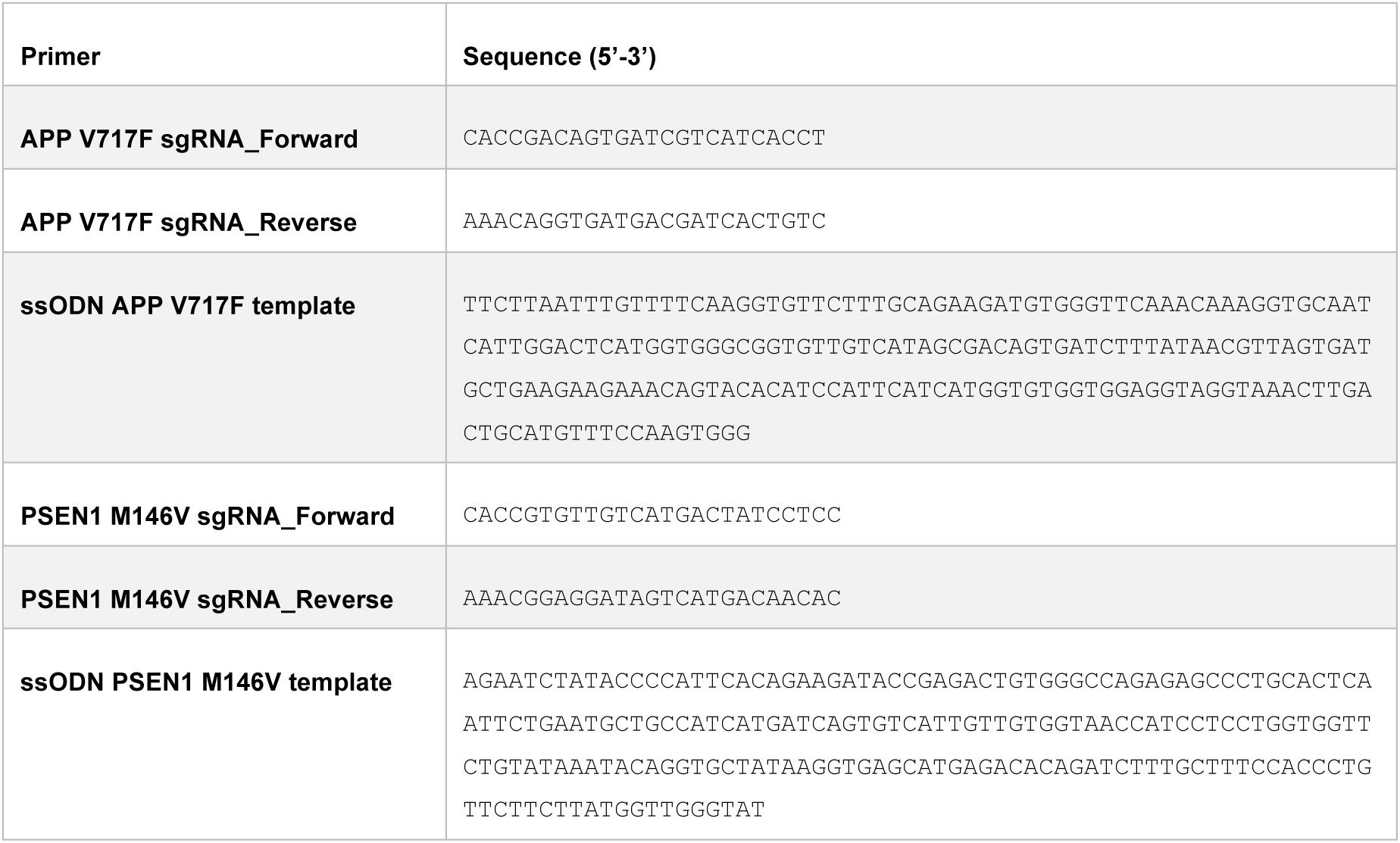
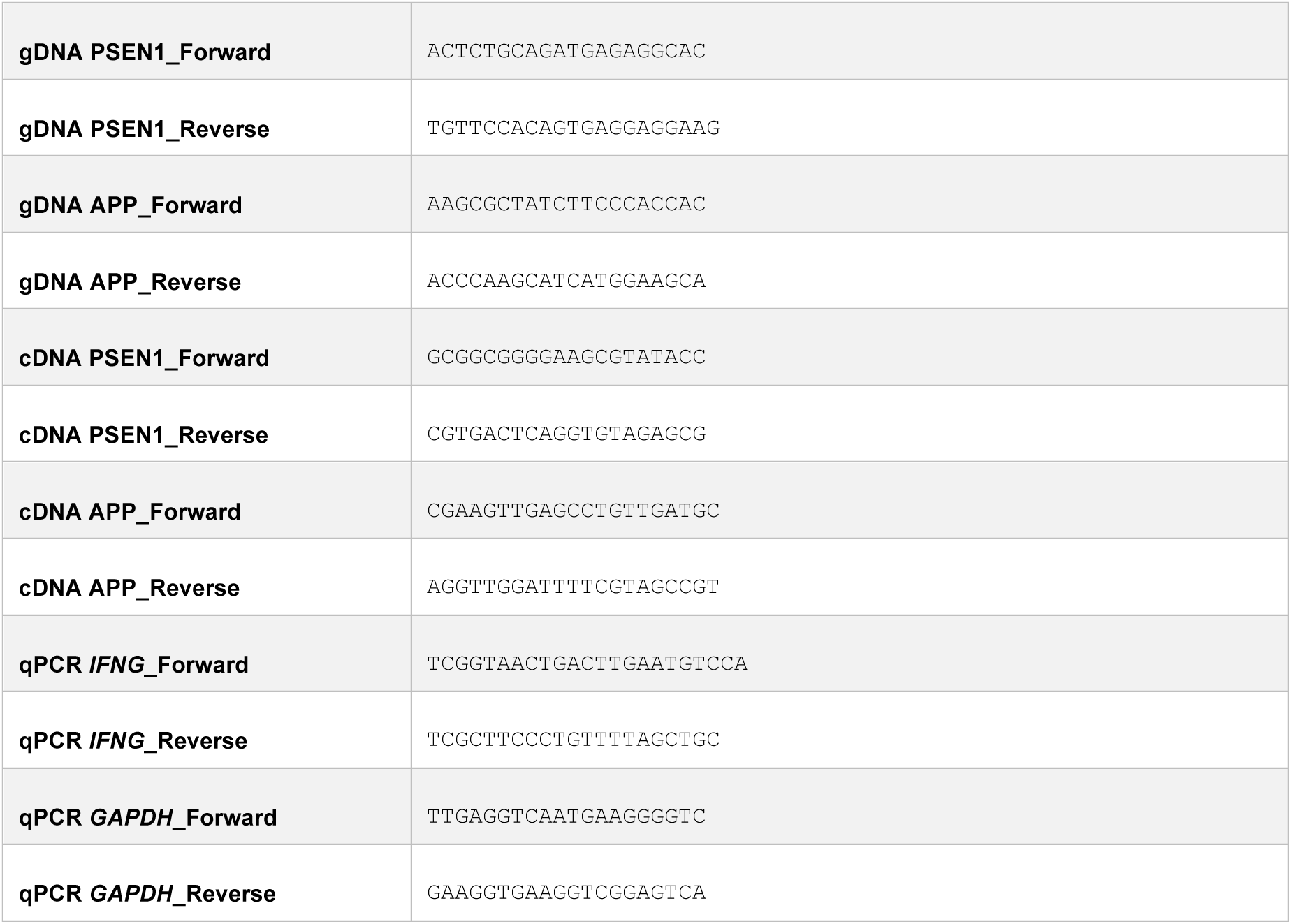
Sequences for CRISPR/Cas9 gene editing and analysis.

## References

1. Wishart, D.S. Chapter 3: Small Molecules and Disease. PLOS Computational Biology 8, e1002805 (2012).

2. Duran-Frigola, M., Rossell, D. & Aloy, P. A chemo-centric view of human health and disease. Nature Communications 5, 5676 (2014).

3. Sterling, T. & Irwin, J.J. ZINC 15 – Ligand Discovery for Everyone. Journal of Chemical Information and Modeling 55, 2324–2337 (2015).

4. Gaulton, A. et al. The ChEMBL database in 2017. Nucleic Acids Res 45, D945–D954 (2017).

5. Wang, Y., et al. PubChem BioAssay: 2017 update. Nucleic Acids Res 45, D955–D963 (2017).

6. Rouillard, A.D. et al. The harmonizome: a collection of processed datasets gathered to serve and mine knowledge about genes and proteins. Database 2016, baw100-baw100 (2016).

7. Newman, D.J. & Cragg, G.M. Natural Products as Sources of New Drugs from 1981 to 2014. Journal of Natural Products 79, 629–661 (2016).

8. Rodrigues, T., Reker, D., Schneider, P. & Schneider, G. Counting on natural products for drug design. Nature Chemistry 8, 531 (2016).

9. Welsch, M.E., Snyder, S.A. & Stockwell, B.R. Privileged scaffolds for library design and drug discovery. Current Opinion in Chemical Biology 14, 347–361 (2010).

10. Bleicher, K.H., Böhm, H.-J., Müller, K. & Alanine, A.I. Hit and lead generation: beyond high-throughput screening. Nature Reviews Drug Discovery 2, 369 (2003).

11. Holbeck, S.L., Collins, J.M. & Doroshow, J.H. Analysis of Food and Drug Administration– Approved Anticancer Agents in the NCI60 Panel of Human Tumor Cell Lines. Molecular Cancer Therapeutics 9, 1451 (2010).

12. Campillos, M., Kuhn, M., Gavin, A.-C., Jensen, L.J. & Bork, P. Drug Target Identification Using Side-Effect Similarity. Science 321, 263 (2008).

13. Petrone, P.M. et al. Rethinking Molecular Similarity: Comparing Compounds on the Basis of Biological Activity. ACS Chemical Biology 7, 1399–1409 (2012).

14. Papadatos, G., Gaulton, A., Hersey, A. & Overington, J.P. Activity, assay and target data curation and quality in the ChEMBL database. J Comput Aided Mol Des 29, 885–896 (2015).

15. Duran-Frigola, M., Mateo, L. & Aloy, P. Drug repositioning beyond the low-hanging fruits. Current Opinion in Systems Biology 3, 95–102 (2017).

16. Nguyen, D.T. et al. Pharos: Collating protein information to shed light on the druggable genome. Nucleic Acids Res 45, D995–D1002 (2017).

17. Duran-Frigola, M., Fernandez-Torras, A., Bertoni, M. & Aloy, P. Formatting biological big data for modern machine learning in drug discovery. WIREs Comp Mol Sci (2018).

18. Corsello, S.M. et al. The Drug Repurposing Hub: a next-generation drug library and information resource. Nat Med 23, 405–408 (2017).

19. Jokinen, E. & Koivunen, J.P. MEK and PI3K inhibition in solid tumors: rationale and evidence to date. Ther Adv Med Oncol 7, 170–180 (2015).

20. Lamb, J. et al. The Connectivity Map: Using Gene-Expression Signatures to Connect Small Molecules, Genes, and Disease. Science 313, 1929 (2006).

21. Subramanian, A. et al. A Next Generation Connectivity Map: L1000 Platform and the First 1,000,000 Profiles. Cell 171, 1437–1452 e1417 (2017).

22. Filzen, T.M., Kutchukian, P.S., Hermes, J.D., Li, J. & Tudor, M. Representing high throughput expression profiles via perturbation barcodes reveals compound targets. PLOS Computational Biology 13, e1005335 (2017).

23. Chen, B. et al. Reversal of cancer gene expression correlates with drug efficacy and reveals therapeutic targets. Nature Communications 8, 16022 (2017).

24. Iorio, F. et al. A Landscape of Pharmacogenomic Interactions in Cancer. Cell 166, 740–754 (2016).

25. Encinas, M. et al. Sequential treatment of SH-SY5Y cells with retinoic acid and brain-derived neurotrophic factor gives rise to fully differentiated, neurotrophic factor-dependent, human neuron-like cells. J Neurochem 75, 991–1003 (2000).

26. Tanzi, R.E. The genetics of Alzheimer disease. Cold Spring Harb Perspect Med 2 (2012).

27. Carvalho-Silva, D. et al. Open Targets Platform: new developments and updates two years on. Nucleic Acids Res 47, D1056–D1065 (2019).

28. Perszyk, R.E. et al. GluN2D-Containing N-methyl-d-Aspartate Receptors Mediate Synaptic Transmission in Hippocampal Interneurons and Regulate Interneuron Activity. Mol Pharmacol 90, 689–702 (2016).

29. Harold, D. et al. Genome-wide association study identifies variants at CLU and PICALM associated with Alzheimer’s disease. Nat Genet 41, 1088–1093 (2009).

30. Anselmo, A.C., Gokarn, Y. & Mitragotri, S. Non-invasive delivery strategies for biologics. Nat Rev Drug Discov 18, 19–40 (2019).

31. Depper, J.M., Leonard, W.J., Robb, R.J., Waldmann, T.A. & Greene, W.C. Blockade of the interleukin-2 receptor by anti-Tac antibody: inhibition of human lymphocyte activation. J Immunol 131, 690–696 (1983).

32. Benson, J.M. et al. Therapeutic targeting of the IL-12/23 pathways: generation and characterization of ustekinumab. Nat Biotechnol 29, 615–624 (2011).

33. Reddy, M., et al. Modulation of CLA, IL-12R, CD40L, and IL-2Ralpha expression and inhibition of IL-12- and IL-23-induced cytokine secretion by CNTO 1275. Cell Immunol 247, 1–11 (2007).

34. Xu, M.J., Johnson, D.E. & Grandis, J.R. EGFR-targeted therapies in the post-genomic era. Cancer Metastasis Rev 36, 463–473 (2017).

35. Masuelli, L. et al. Apigenin induces apoptosis and impairs head and neck carcinomas EGFR/ErbB2 signaling. Front Biosci (Landmark Ed) 16, 1060–1068 (2011).

36. Hu, W.J., Liu, J., Zhong, L.K. & Wang, J. Apigenin enhances the antitumor effects of cetuximab in nasopharyngeal carcinoma by inhibiting EGFR signaling. Biomed Pharmacother 102, 681–688 (2018).

37. Sawai, A. et al. Inhibition of Hsp90 down-regulates mutant epidermal growth factor receptor (EGFR) expression and sensitizes EGFR mutant tumors to paclitaxel. Cancer Res 68, 589–596 (2008).

38. Williams, A.J. et al. Open PHACTS: semantic interoperability for drug discovery. Drug Discovery Today 17, 1188–1198 (2012).

39. Rodgers, G. et al. Glimmers in illuminating the druggable genome. Nature Reviews Drug Discovery 17, 301 (2018).

40. Reymond, J.-L. The Chemical Space Project. Accounts of Chemical Research 48, 722–730 (2015).

41. Irwin, J.J., Gaskins, G., Sterling, T., Mysinger, M.M. & Keiser, M.J. Predicted Biological Activity of Purchasable Chemical Space. Journal of Chemical Information and Modeling 58, 148–164 (2018).

42. Axen, S.D. et al. A Simple Representation of Three-Dimensional Molecular Structure. J Med Chem 60, 7393–7409 (2017).

43. Bemis, G.W. & Murcko, M.A. The properties of known drugs. 1. Molecular frameworks. J Med Chem 39, 2887–2893 (1996).

44. Durant, J.L., Leland, B.A., Henry, D.R. & Nourse, J.G. Reoptimization of MDL keys for use in drug discovery. J Chem Inf Comput Sci 42, 1273–1280 (2002).

45. Bickerton, G.R., Paolini, G.V., Besnard, J., Muresan, S. & Hopkins, A.L. Quantifying the chemical beauty of drugs. Nat Chem 4, 90–98 (2012).

46. Lipinski, C.A. Lead- and drug-like compounds: the rule-of-five revolution. Drug Discov Today Technol 1, 337–341 (2004).

47. Congreve, M., Carr, R., Murray, C. & Jhoti, H. A ’rule of three’ for fragment-based lead discovery? Drug Discov Today 8, 876–877 (2003).

48. Wishart, D.S. et al. DrugBank 5.0: a major update to the DrugBank database for 2018. Nucleic Acids Res 46, D1074–D1082 (2018).

49. Cheng, H. et al. ECOD: an evolutionary classification of protein domains. PLoS Comput Biol 10, e1003926 (2014).

50. Gilson, M.K., et al. BindingDB in 2015: A public database for medicinal chemistry, computational chemistry and systems pharmacology. Nucleic Acids Res 44, D1045–1053 (2016).

51. Hastings, J., et al. ChEBI in 2016: Improved services and an expanding collection of metabolites. Nucleic Acids Res 44, D1214–1219 (2016).

52. Thiele, I. et al. A community-driven global reconstruction of human metabolism. Nat Biotechnol 31, 419–425 (2013).

53. Cerami, E.G. et al. Pathway Commons, a web resource for biological pathway data. Nucleic Acids Res 39, D685–690 (2011).

54. Fabregat, A. et al. The Reactome Pathway Knowledgebase. Nucleic Acids Res 46, D649–D655 (2018).

55. Pryszcz, L.P., Huerta-Cepas, J. & Gabaldon, T. MetaPhOrs: orthology and paralogy predictions from multiple phylogenetic evidence using a consistency-based confidence score. Nucleic Acids Res 39, e32 (2011).

56. Kruger, F.A. & Overington, J.P. Global analysis of small molecule binding to related protein targets. PLoS Comput Biol 8, e1002333 (2012).

57. Zwierzyna, M. & Overington, J.P. Classification and analysis of a large collection of in vivo bioassay descriptions. PLOS Computational Biology 13, e1005641 (2017).

58. Szklarczyk, D. et al. The STRING database in 2017: quality-controlled protein-protein association networks, made broadly accessible. Nucleic Acids Res 45, D362–D368 (2017).

59. Li, T. et al. A scored human protein-protein interaction network to catalyze genomic interpretation. Nat Methods 14, 61–64 (2017).

60. Kanehisa, M., Sato, Y., Kawashima, M., Furumichi, M. & Tanabe, M. KEGG as a reference resource for gene and protein annotation. Nucleic Acids Res 44, D457–462 (2016).

61. Kandasamy, K. et al. NetPath: a public resource of curated signal transduction pathways. Genome Biol 11, R3 (2010).

62. Mi, H. et al. PANTHER version 11: expanded annotation data from Gene Ontology and Reactome pathways, and data analysis tool enhancements. Nucleic Acids Res 45, D183–D189 (2017).

63. Kelder, T. et al. WikiPathways: building research communities on biological pathways. Nucleic Acids Res 40, D1301–1307 (2012).

64. Mosca, R., Ceol, A. & Aloy, P. Interactome3D: adding structural details to protein networks. Nat Methods 10, 47–53 (2013).

65. Leiserson, M.D. et al. Pan-cancer network analysis identifies combinations of rare somatic mutations across pathways and protein complexes. Nat Genet 47, 106–114 (2015).

66. Iorio, F. et al. Discovery of drug mode of action and drug repositioning from transcriptional responses. Proc Natl Acad Sci U S A 107, 14621–14626 (2010).

67. Barretina, J. et al. The Cancer Cell Line Encyclopedia enables predictive modelling of anticancer drug sensitivity. Nature 483, 603–607 (2012).

68. Basu, A. et al. An interactive resource to identify cancer genetic and lineage dependencies targeted by small molecules. Cell 154, 1151–1161 (2013).

69. Chabner, B.A. NCI-60 Cell Line Screening: A Radical Departure in its Time. J Natl Cancer Inst 108 (2016).

70. Azur, M.J., Stuart, E.A., Frangakis, C. & Leaf, P.J. Multiple imputation by chained equations: what is it and how does it work? Int J Methods Psychiatr Res 20, 40–49 (2011).

71. Nelson, J. et al. MOSAIC: a chemical-genetic interaction data repository and web resource for exploring chemical modes of action. Bioinformatics (2017).

72. Wawer, M.J. et al. Toward performance-diverse small-molecule libraries for cell-based phenotypic screening using multiplexed high-dimensional profiling. Proc Natl Acad Sci U S A 111, 10911–10916 (2014).

73. Brown, A.S. & Patel, C.J. A standard database for drug repositioning. Sci Data 4, 170029 (2017).

74. Piñero, J. et al. DisGeNET: a comprehensive platform integrating information on human disease-associated genes and variants. Nucleic Acids Research 45, D833–D839 (2017).

75. Kuhn, M., Letunic, I., Jensen, L.J. & Bork, P. The SIDER database of drugs and side effects. Nucleic Acids Res 44, D1075–1079 (2016).

76. Kuhn, M. et al. Systematic identification of proteins that elicit drug side effects. Mol Syst Biol 9, 663 (2013).

77. Duran-Frigola, M. & Aloy, P. Analysis of chemical and biological features yields mechanistic insights into drug side effects. Chem Biol 20, 594–603 (2013).

78. Davis, A.P. et al. The Comparative Toxicogenomics Database: update 2017. Nucleic Acids Res 45, D972–D978 (2017).

79. Ryu, J.Y., Kim, H.U. & Lee, S.Y. Deep learning improves prediction of drug–drug and drug– food interactions. Proceedings of the National Academy of Sciences 115, E4304–E4311 (2018).

80. Grover, A. & Leskovec, J. node2vec: Scalable Feature Learning for Networks. arXiv:1607.00653 (2016).

81. Matsui, Y.O., K; Yamasaki, T; Aizawa K PQk-means: Billion-scale Clustering for Product-quantized Codes. arXiv:1709.03708 (2017).

82. Maaten, L.v.d. Barnes-Hut-SNE. arXiv:1301.3342 (2013).

83. 83. McInnes, L. & Healy, J. in 2017 IEEE International Conference on Data Mining Workshops (ICDMW) (2017).

84. Webber, W., Moffat, A. & Zobel, J. A similarity measure for indefinite rankings. ACM Trans. Inf. Syst. 28, 1–38 (2010).

85. Lo, Y.C. et al. Large-scale chemical similarity networks for target profiling of compounds identified in cell-based chemical screens. PLoS Comput Biol 11, e1004153 (2015).

86. Rennie, J.D.M., Shih, L., Teevan, J. & Karger, D.R. in International Conference on International Conference on Machine Learning 616–623 (AAAI Press, Washington, DC, USA; 2003).

87. Irwin, J.J. & Shoichet, B.K. ZINC--a free database of commercially available compounds for virtual screening. J Chem Inf Model 45, 177–182 (2005).

88. Fernandez-Torras, A., Duran-Frigola, M. & Aloy, P. Encircling the regions of the pharmacogenomic landscape that determine drug response. Genome Medicine 26, 17 (2019).

89. Badia, R. et al. SAMHD1 is active in cycling cells permissive to HIV-1 infection. Antiviral Res 142, 123–135 (2017).

90. Saxena, V., Orgill, D. & Kohane, I. Absolute enrichment: gene set enrichment analysis for homeostatic systems. Nucleic Acids Res 34, e151 (2006).

